# Structural basis for tunable control of actin dynamics by myosin-15 in mechanosensory stereocilia

**DOI:** 10.1101/2021.07.09.451843

**Authors:** Rui Gong, Fangfang Jiang, Zane G. Moreland, Matthew J. Reynolds, Santiago Espinosa de los Reyes, Pinar S. Gurel, Arik Shams, Michael R. Bowl, Jonathan E. Bird, Gregory M. Alushin

## Abstract

The motor protein myosin-15 is necessary for the development and maintenance of mechanosensory stereocilia, and myosin-15 mutations cause profound deafness. In a companion study, we report that myosin-15 nucleates actin filament (“F-actin”) assembly and identify a progressive hearing loss mutation (p.D1647G, “*jordan*”) which disrupts stereocilia elongation by inhibiting actin polymerization. Here, we present cryo-EM structures of myosin-15 bound to F-actin, providing a framework for interpreting deafness mutations and their impacts on myosin-stimulated actin assembly. Rigor myosin-15 evokes conformational changes in F-actin yet maintains flexibility in actin’s D-loop, which mediates inter-subunit contacts, while the *jordan* mutant locks the D-loop in a single conformation. ADP-bound myosin-15 also locks the D-loop, which correspondingly blunts actin-polymerization stimulation. We propose myosin-15 enhances polymerization by bridging actin protomers, regulating nucleation efficiency by modulating actin’s structural plasticity in a myosin nucleotide-state dependent manner. This tunable regulation of actin polymerization could be harnessed to precisely control stereocilium height.

## Introduction

The stereocilia of inner ear hair cells are rod-like membrane protrusions responsible for sound detection. Each hair cell assembles a staircase-shaped mechanosensory hair bundle during development, which contains three rows of stereocilia of increasing height. Stereocilia are primarily composed of parallel bundles of filamentous actin (F-actin) crosslinked to form a para-crystalline array (DeRosier et al., 1980; Tilney et al., 1980). Stereocilia develop from microvilli that undergo an exquisite morphogenetic program characterized by F-actin elongation and bundling, presumed to be orchestrated by coordinated actin polymerization and bundling protein engagement (Tilney et al., 1992). Once a stereocilium is established, slow actin dynamics maintain its architecture throughout adult life (Drummond et al., 2015; Narayanan et al., 2015; Zhang et al., 2012). Finely-tuned regulation of actin dynamics is critical for stereocilium height determination, and mutations in actin and its regulatory factors are frequently associated with hearing loss (Barr-Gillespie, 2015; Drummond et al., 2012). Nevertheless, the molecular mechanisms which precisely define the height of individual stereocilia during development and maintain them for a lifetime remain elusive.

Unconventional myosin-15 (encoded by the *Myo15* gene in mouse, and *MYO15A* in humans) is a plus-end directed molecular motor essential for stereocilia development and maintenance (Anderson et al., 2000; Belyantseva et al., 2003; Belyantseva et al., 2005; Bird et al., 2014; Fang et al., 2015; Probst et al., 1998; Wang et al., 1998). Over 300 mutations have been mapped to the human *MYO15A* gene that cause autosomal recessive non-syndromic deafness, DNFB3 (Rehman et al., 2016; Wang et al., 1998). Within hair cells, myosin-15 localizes to the distal tips of stereocilia, the major sites of actin turnover which are enriched with F-actin plus (“barbed”) ends (Belyantseva et al., 2003; Belyantseva et al., 2005; Drummond et al., 2015; Narayanan et al., 2015; Rzadzinska et al., 2004). In mice homozygous for the *Myo15* “*shaker 2*” allele (p.C1779Y), myosin-15 trafficking is disrupted in nascent stereocilia (Belyantseva 2003), resulting in failure of stereocilia to elongate and profound hearing loss (Probst et al., 1998). Myosin-15 transports whirlin (WHRN), EPS8, GPSM2, and GNAI3 to stereocilia tips, an “elongation network” of interacting proteins required for stereocilium biogenesis, each member of which phenocopies *shaker 2* when individually mutated in mice (Belyantseva et al., 2005; Delprat et al., 2005; Manor et al., 2011; Mauriac et al., 2017; Mburu et al., 2003; Tadenev et al., 2019; Tarchini et al., 2016; Zampini et al., 2011). As EPS8 is a known actin-binding protein (ABP) which exhibits filament bundling and plus-end capping activity (Disanza et al., 2004; Hertzog et al., 2010), it has been proposed that myosin-15 mediates stereocilia growth by transporting elongation network components to the tip compartment, where they serve as the regulatory machinery governing F-actin assembly (Belyantseva et al., 2005; Delprat et al., 2005; Manor et al., 2011; Mauriac et al., 2017; Tadenev et al., 2019). Nevertheless, the functional consequences of specific myosin-15 mutations remain poorly understood due to the absence of structural data and limited functional characterization of the myosin-15 protein.

In a companion manuscript, we identified the myosin-15 missense mutation p.D1647G, hereafter referred to as the “*jordan*” mutant (Figure 1A), in a forward genetic screen for lesions leading to progressive hearing loss in mice (Moreland, 2021). The *jordan* mouse develops abnormally short stereocilia which retract during aging. Although its actin-activated ATPase activity and F-actin binding affinity are somewhat compromised, purified *jordan* myosin-15 is nevertheless an active motor, and WHRN, EPS8, GPSM2 and GNAI3 localize to the tips of developing stereocilia in the *jordan* mouse, leading us to hypothesize that myosin-15 could control stereocilium height through additional mechanisms. Consistently, we found that the motor-domain containing myosin-15 S1 fragment dramatically stimulates actin polymerization *in vitro*, with the nucleotide-free “rigor” state specifically acting as a nucleation factor, while the *jordan* mutant instead suppresses actin assembly (Moreland, 2021). Early studies of the S1 fragment of skeletal muscle myosin demonstrated its capacity to nucleate actin polymerization *in vitro* (Fievez et al., 1997; Lheureux et al., 1993; Miller et al., 1988), but a physiological role for this activity has not, to our knowledge, been reported. Our studies suggest myosin-15 directly regulates actin dynamics in stereocilia, but the underlying molecular mechanism, and how it is disrupted by a single amino acid substitution in the *jordan* mutant, remain unknown.

**Figure 1.**
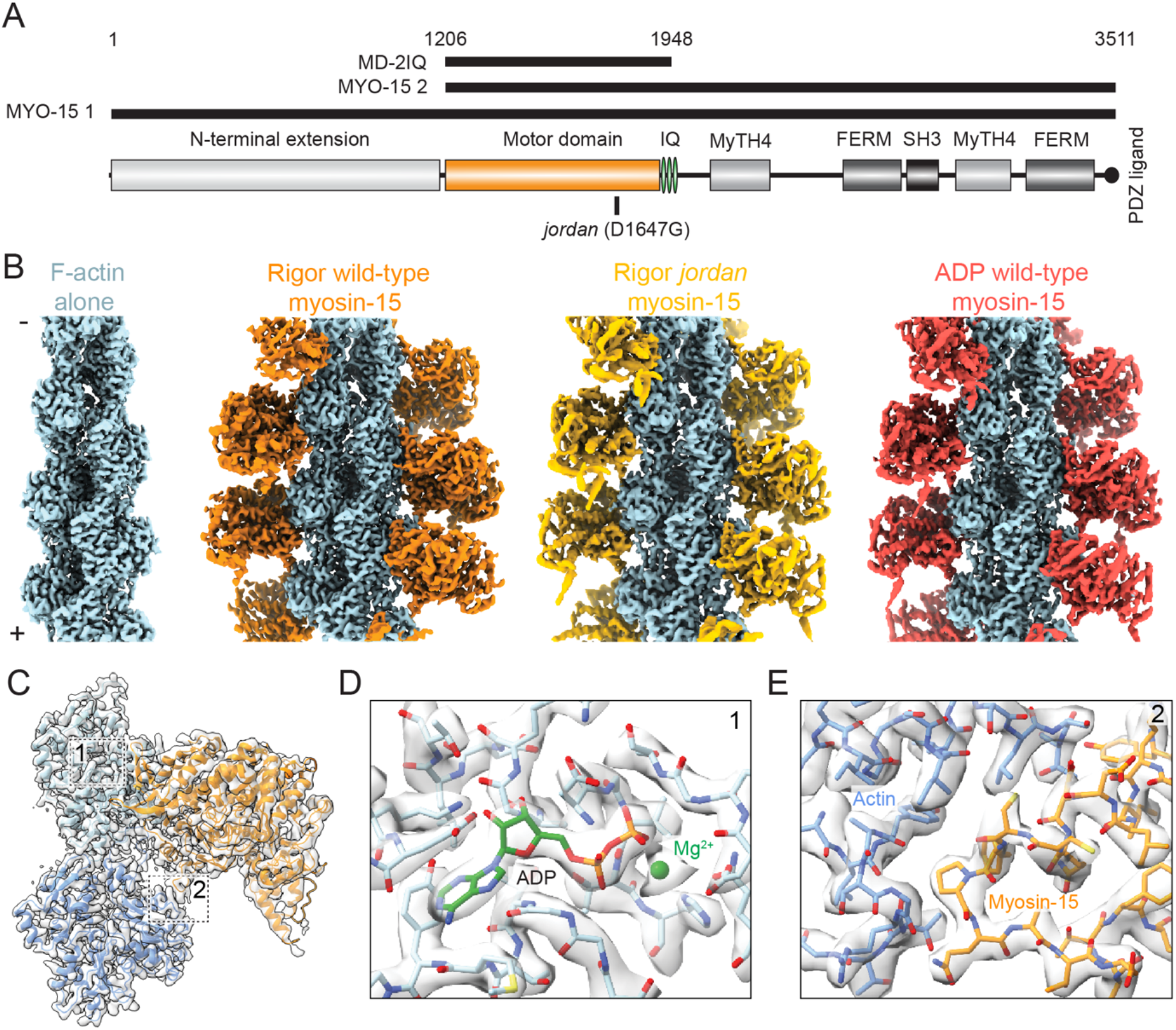
Cryo-EM structures of actomyosin-15 complexes. **(A)** Diagram of *M. musculus* myosin-15 primary structure. **(B)** Cryo-EM maps of F-actin alone and indicated actomyosin-15 complexes. **(C)** Cryo-EM map and ribbon diagram of two actin and one myosin-15 subunits from the rigor wild-type actomyosin-15 reconstruction. Numbered boxes correspond to detail views in subsequent panels. **(D)** Cryo-EM map and stick models around actin’s ADP binding pocket. **(E)** Cryo-EM map and stick models at the actin-myosin loop 3 interface.

F-actin assembly and disassembly are tightly regulated by dozens of ABPs, including severing proteins, capping proteins, and nucleators, whose activities must be precisely coordinated to construct elaborated cellular protrusions such as stereocilia (Pollard, 2016; Tilney et al., 1992). These ABPs frequently utilize their actin-binding interfaces to modulate G-actin monomer / F-actin protomer conformation and thereby fine-tune actin polymerization dynamics (Crevenna et al., 2015; Dominguez and Holmes, 2011; Pollard, 2016). The structurally polymorphic actin subdomain 2, or “D-loop”, which mediates longitudinal interactions between F-actin protomers, plays a prominent role in this conformational regulation (Das et al., 2020; Durer et al., 2012; Galkin et al., 2010). Several myosin motors have been reported to modulate actin’s conformation through direct contacts with the D-loop upon F-actin binding (Gurel et al., 2017; Mentes et al., 2018; von der Ecken et al., 2016), and distinct actin conformations were visualized at the Mg-ADP and rigor stages of myosin-6’s mechanochemical cycle (Gurel et al., 2017). This led us to hypothesize that reciprocal conformational changes at the actin-myosin interface could be harnessed by myosin-15 during its mechanochemical cycle to precisely regulate F-actin polymerization dynamics to control stereocilium height.

Here we report the cryo-electron microscopy (cryo-EM) structure of the rigor wild-type myosin-15 motor domain bound to F-actin at 2.84 Å resolution, to our knowledge the highest-resolution actomyosin structure to date, providing a framework for broadly interpreting deafness mutations in the motor domain and at the actin-myosin interface. To probe the mechanisms by which myosin-15 regulates F-actin assembly, we also obtained cryo-EM structures of F-actin alone (2.82 Å), rigor *jordan* myosin-15 (3.76 Å), and Mg-ADP wild-type myosin-15 (3.63 Å) bound to F-actin. The unprecedented resolution of our F-actin alone structure reveals a region of the D-loop (residues G48-Q49) that is structurally flexible but nevertheless primarily adopts a mixture of two conformations. Rigor wild-type myosin-15 engagement elicits rearrangements at the actin-myosin interface which are propagated through the actin structure, yet the D-loop continues to adopt two conformations. Binding of the *jordan* mutant evokes similar overall rearrangements but strikingly locks the D-loop in a single conformation, as does Mg-ADP wild-type myosin-15, which shows reduced F-actin stimulation relative to the rigor state in polymerization assays. Our studies suggest strongly-bound myosin-15 promotes F-actin assembly by reinforcing longitudinal contacts between actin protomers with a nucleation efficiency controlled by the motor domain’s nucleotide-state, which modulates actin conformational flexibility mediating F-actin polymerization. We propose this tunable direct regulation of actin assembly dynamics facilitates precise control of stereocilia height by myosin-15 *in vivo*, which is completely compromised by the *jordan* deafness mutation.

## Results

### Cryo-EM reconstructions of actomyosin-15 complexes and bare F-actin

Using cryo-EM and the iterative-real space helical symmetry reconstruction (IHRSR) procedure as implemented in RELION-3 (Egelman, 2007; Zivanov et al., 2018) we determined structures of a wild-type mouse myosin-15 construct containing the motor domain and 2 IQ motifs (Bird et al., 2014) (“MD-2IQ”, Figure 1A) bound to F-actin at a 1:1 stoichiometry in the rigor and Mg-ADP states at an average resolution of 2.84 Å and 3.63 Å, respectively (STAR Methods). We additionally determined the structure of the *jordan* deafness mutant bound to F-actin at 3.76 Å resolution (Figures 1B, S1A,B). Our rigor wild-type structure represents a substantial improvement in resolution over previously reported actomyosin reconstructions (Gurel et al., 2017; Mentes et al., 2018; Risi et al., 2021; von der Ecken et al., 2016), facilitating building an accurate atomic model of the myosin-15 motor domain (Figure 1C), which has not, to our knowledge, previously been structurally visualized in the absence or presence of F-actin.

As has been reported in other cryo-EM studies of actomyosin (Gurel et al., 2017; Mentes et al., 2018; Risi et al., 2021; von der Ecken et al., 2016), the local resolution of these reconstructions is highest in the center of the filament and decreases radially outwards towards the distal tip of the myosin lever arm (Figure S1B). For all three complexes, the resolution of F-actin and the actin-myosin interface extends beyond 3 Å, yielding models with unambiguous side-chain density within these areas (Figures 1D,E; S1B). Most side chains in the motor domain are clearly resolved and secondary structures can be identified in and around the distal converter domain (Figures 1E, S1B). To visualize structural changes evoked in F-actin by myosin-15, we also reconstructed bare F-actin at a global resolution of 2.82 Å (Figures 1B; S1A,B), to our knowledge the highest resolution mammalian F-actin structure reported to date. This density map facilitated direct model building and refinement of actin’s bound Mg-ADP and all actin residues except the first five at the protein’s flexible N terminus.

### Structural interpretation of deafness-causing myosin-15 mutations

Within the motor domain of myosin-15, a total of 59 missense mutations targeting 58 residues have been identified in DFNB3 patients (Rehman et al., 2016; Zhang et al., 2019). Of these 58 residues, 2 are located in disordered regions which were not resolved in our reconstruction: the remaining 56 can be mapped onto the structure (Figure 2A). Sequence alignment of the mouse and human myosin-15 motor domains shows 54 of the 56 residues are identical and 2 of them are highly similar (Figure S2). Examining the distribution of these deafness-causing mutations leads us to classify them into four groups (Figure 2A: unless otherwise noted, residues cited correspond to the human protein). Mutations in the first group are found in the core of the motor domain and are likely to disrupt proper folding and thereby completely compromise function. For example, the p.S1465P mutation introduces a proline in the middle of a long helix and the p.A1532T/p.A1535D mutations substitute alanine with a polar or charged amino acid in the hydrophobic core (Figure 2B), all of which we anticipate will severely impede folding.

**Figure 2.**
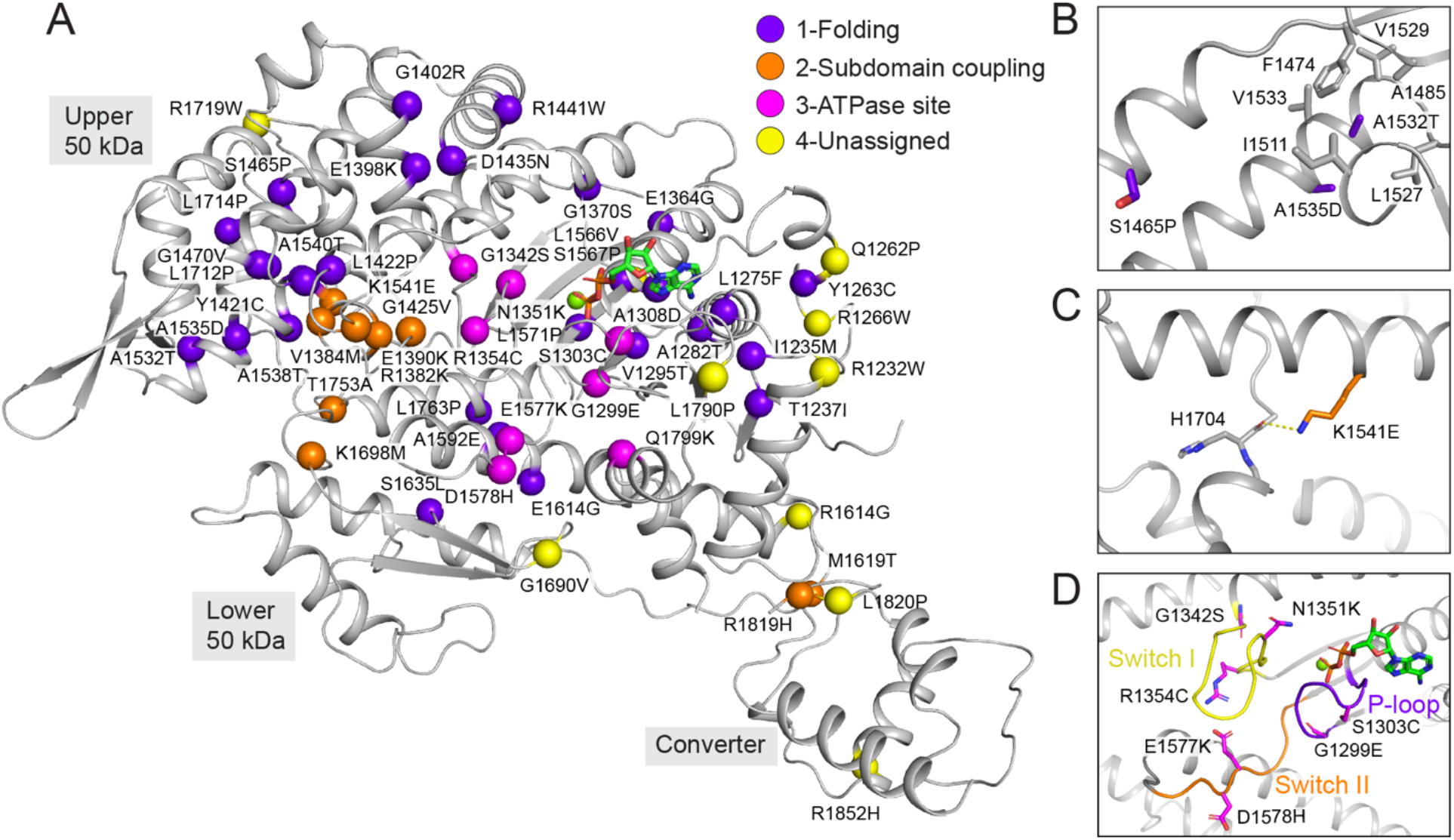
Structural mapping of deafness causing myosin-15 mutations. **(A)** Ribbon diagram of the myosin-15 motor domain with *α*-carbons at sites of deafness causing mutations shown as spheres, colored according to the indicated mechanistic categories. **(B)** Detail view of mutations anticipated to disrupt folding. **(C)** Detail view of mutation anticipated to affect subdomain rearrangements. **(D)** Detail view of mutations in the switch I, II and P-loop regions anticipated to impact ATPase activity.

This group additionally includes mouse C1779, the site of the pathogenic p.C1779Y deafness mutation in the *shaker 2* allele, contrary to its originally predicted location at an actin-binding interface (Probst et al., 1998). This residue sits in a hydrophobic core surrounded by mouse I1297, L1275, A1308 and V1777 (Figure S3A), where substitution with a bulky hydrophilic tyrosine would be anticipated to disrupt the proper folding of the transducer. Consistent with this interpretation, recombinantly expressed *shaker 2* (p.C1779Y) motor domain is a soluble aggregate when purified and subjected to size exclusion chromatography (Moreland, 2021). Furthermore, the human counterpart (A1324) of mouse A1308, which is adjacent to C1779 (Figure S3A), was also found to be substituted with the highly polar residue aspartic acid in a pedigree with profound deafness (Zhang et al., 2018), consistent with disruption of the hydrophobic core resulting in misfolding and aggregation.

Mutations in the second group are located in regions mediating interactions between the motor’s subdomains, which are likely to interfere with coordinated rearrangements required to execute the ATPase mechanochemical cycle (Houdusse and Sweeney, 2016) The side chain of the highly conserved K1541 from the upper 50-kDa subdomain forms a hydrogen bond with the backbone of H1704 in the loop connecting the upper 50-kDa and lower 50-kDa subdomains (Figure 2C and Figure S2). This interaction likely plays a stabilizing role in the cleft, and the charge inversion p.K1541E mutation is anticipated to disrupt it and hinder cleft closure. The third group are located in consensus ATPase motifs, including switch I, switch II, and the P-loop. These mutations are expected to have a deleterious impact on the mechanochemical cycle by impeding ATP binding and/or hydrolysis (Figure 2D and Figure S2). Finally, a fourth group of mutations are distributed in solvent exposed loop regions with no obvious pattern (Figure 2A). We speculate these residues mediate binding contacts with yet to be identified partner proteins or intramolecular contacts involved in regulating myosin-15’s motor function, which could include an interface between the motor domain and the large 133 kDa N-terminal domain of myosin-15 isoform 1 (MYO15-1; Figure 1A) that is required for postnatal stereocilia maintenance (Fang et al., 2015). Our structural mapping analysis suggests that mutations in myosin-15 cause pathology by either interfering with canonical myosin-15 ATPase motor activity, or with non-canonical functions such as mediating protein-protein interactions at the stereocilia tips.

### A structurally diversified myosin-15-actin interface

We next undertook a detailed analysis of the myosin-15-F-actin interface to identify specific contacts which could contribute to myosin-15’s F-actin nucleation activity. We also examined the interface for contacts which might be impacted by deafness mutations in both myosin-15 and *γ*-actin (encoded by the ACTG1 gene in humans), one of the primary actin isoforms expressed in hair cells (Figure S3B). Like other myosins, myosin-15 employs conserved structural elements to interact with F-actin: the Helix-Loop-Helix (HLH) motif, the cardiomyopathy (CM) loop, loop 2, loop 3, and the activation loop (Figure 3A and S4A). However, residue-level interactions have diversified among myosins to fine-tune the functions of superfamily members (Gurel et al., 2017; Mentes et al., 2018; von der Ecken et al., 2016). Myosin-15’s HLH motif (I1641-P1671) mediates the major contact with F-actin through its engagement with two longitudinally adjacent actin subunits (which we here refer to as subunits “i” and “i + 2”) along a protofilament (Figure 3A). The first helix of the HLH includes myosin-15 residue D1647, the site of the p.D1647G mutation in the *jordan* allele, which forms hydrogen bonds with S350 and T351 on subdomain 1 of subunit i (Figure 3B). Notably, D1647 is highly conserved among myosin supefamily members, indicating this interaction is likely broadly important for myosin function (Figure S2). Mutation of this site in myosin-2 resulted in a ten-fold reduction of F-actin binding affinity (Furch et al., 2000), and the F-actin binding of *jordan* mutant myosin-15 is also substantially decreased (Moreland, 2021).

**Figure 3.**
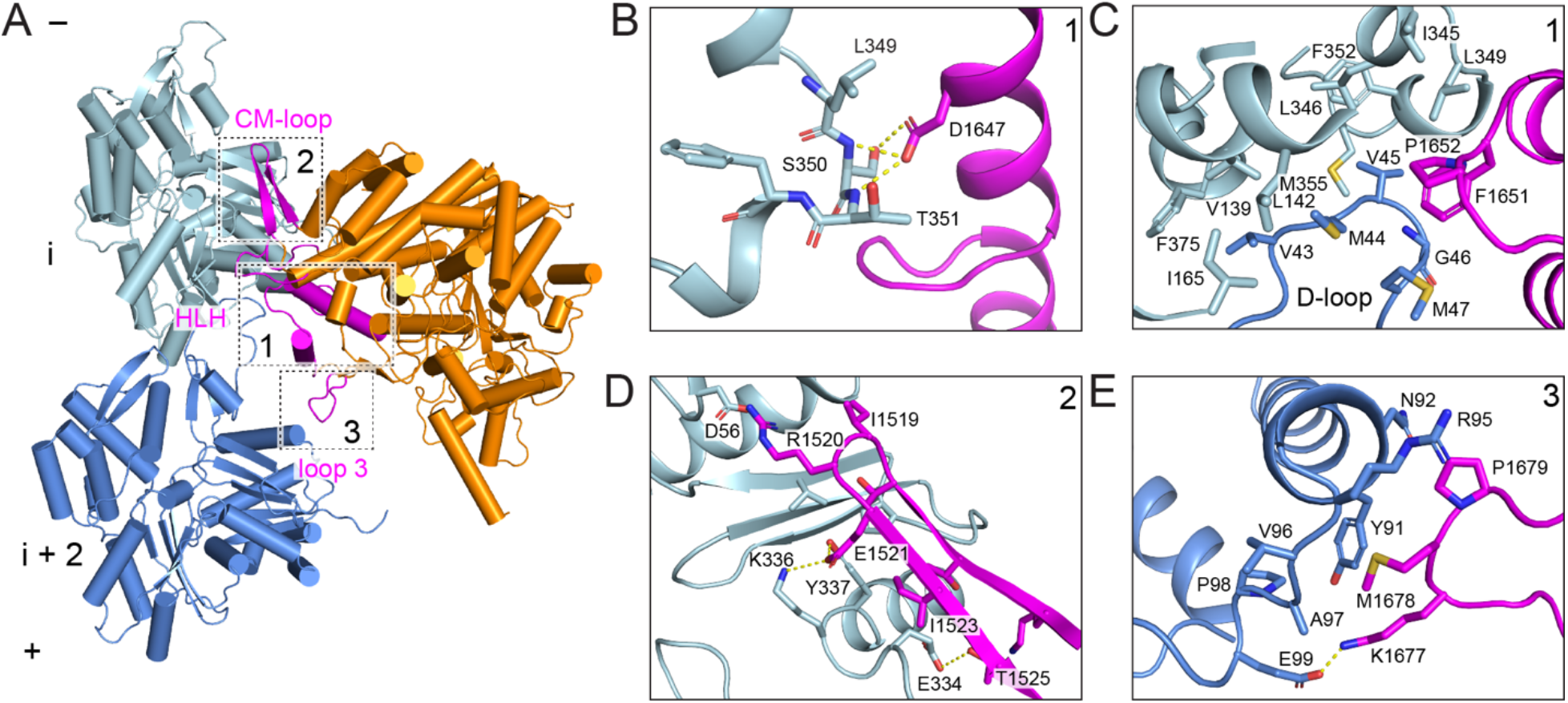
Interactions at the rigor wild-type actomyosin-15 interface. **(A)** Atomic model of the interface between two actin subunits and rigor wild-type myosin-15. Actin subunits i, i+2, and myosin-15 are colored in light blue, cornflower blue, and dark orange, respectively. The indicated myosin-15 actin interaction motifs are highlighted in magenta, and numbered boxes correspond to detail views in subsequent panels. **(B)** Hydrogen bonding interactions between myosin-15 residue D1647 and actin residues S352 and T353. **(C)** Hydrophobic interactions formed between the D-loop of actin subunit i+2, subdomain 1 and 3 of actin subunit i, and the helix-loop-helix (HLH) motif of myosin-15. **(D)** Interface between the CM loop of myosin-15 and subdomain 3 of actin subunit i. **(E)** Contacts between loop 3 of myosin-15 and subdomain 1 of actin subunit i+2.

In the loop portion of the HLH, a pair of hydrophobic residues (F1651 and P1652) inserts into a hydrophobic pocket formed by subdomains 1 and 3 of actin subunit i and the D-loop of subunit i + 2 (Figure 3C), where L349 of subunit i closely engages with myosin-15 F1651. The *γ*-actin mutation L349M causes profound deafness in human patients (Sloan-Heggen et al., 2016), and we anticipate replacement of leucine by the bulkier methionine impairs myosin-15 engagement by preventing access to this pocket (Figure 3C and S3B). Additionally, Q1653 forms a hydrogen bond with the backbone of subunit i residue G146 (Figure S4B). The second helix engages with the N-terminal portion of subunit i + 2’s D-loop through a network of electrostatic interactions. Residue K1662 of myosin-15 forms a hydrogen bond with the sidechain of D-loop residue Q49, and myosin-15 Y1665 also forms a possible N-H···π bond with this actin residue (Figure S4B).

The activation loop (I1634-G1640) preceding the HLH motif has been suggested to activate myosins through contacts with actin’s negatively charged N terminus (Varkuti et al., 2012; Varkuti et al., 2015). Indeed, K1637 forms a possible salt bridge with subunit i + 2 residue 4E, partially stabilizing actin’s flexible N terminus (Figure S4C). This ionic interaction is present in both the Mg-ADP and rigor states (Figure S4C), indicating that it is likely involved in the initial weak-to-strong F-actin binding transition in the case of myosin-15. Loop 2 has also been implicated in mediating initial contacts during the weakly-bound phase of the myosin mechanochemical cycle prior to phosphate release (Murphy and Spudich, 1999; Onishi et al., 2006; Uyeda et al., 1994). This region is largely disordered in our structures, suggesting loop 2 engagement is dispensable for strong binding of myosin-15 to F-actin. Discontinuous density is present for several residues extending from the C-terminal base of loop 2, which orients towards the N terminus of actin and may form weak interactions (Figure S4D). This is in striking contrast to non-muscle myosin-1c (hereafter “myosin-1”) or non-muscle myosin-2c (hereafter “myosin-2”), where loop 2 is either partially or fully ordered at the strongly-bound interface (Mentes et al., 2018; von der Ecken et al., 2016).

The CM loop adopts a beta hairpin fold and packs tightly against subdomain 1 of subunit i (Figure 3A and 3D). In contrast to the CM loops of myosin-1 and myosin-2, which primarily engage in hydrophobic interactions with F-actin (Mentes et al., 2018; von der Ecken et al., 2016), myosin-15’s CM loop-F-actin hydrophobic interface is minimal and is instead dominated by a network of electrostatic interactions. Specifically, the non-conserved residue R1520 at the tip of the CM loop likely mediates a myosin-15-specific salt bridge with actin D56. Similar to myosin-1, the highly conserved residue E1521 mediates electrostatic contacts with actin residues K336 and Y337. The side chain of T1525 also forms a hydrogen bond with actin E334, which is not observed for myosin-1 or myosin-2 (Figure 3D). Myosin-15 possesses a relatively short loop 3 which nevertheless mediates substantial contacts with subdomain 1 of subunit i + 2. Compared with myosin-1 and myosin-2, myosin-15’s loop 3 assumes a distinct conformation to form divergent interactions (Figure S4E). Notably, myosin-15 P1679 makes Van der Waals contacts with actin N92 and R95, and M1678 sits in a shallow hydrophobic groove cradled by actin Y91, V96, A97 and P98. This interface is further strengthened by a probable salt bridge between myosin K1677 and actin E99 (Figure 3E). Collectively, we observe numerous myosin-15-specific interactions with F-actin, consistent with specialization of this interface for myosin-15 function in stereocilia.

### Rigor myosin-15 evokes structural changes in F-actin while maintaining D-loop flexibility

As rigor myosin-15 potently stimulates F-actin polymerization (Moreland, 2021), we next compared two longitudinally adjacent actin subunits from our F-actin alone and rigor wild-type actomyosin-15 structures, hypothesizing myosin-15 could regulate actin polymerization by modulating F-actin conformation. Consistent with previously reported actomyosin structures (Gurel et al., 2017; Mentes et al., 2018; Risi et al., 2021; von der Ecken et al., 2016), the overall conformation of actin in the presence and absence of myosin-15 is highly similar, with a global RMSD of 0.51 Å (Figure 4A). We primarily observe remodeling adjacent to the myosin-15 HLH, which is propagated through the structure into a global compression of the filament along the longitudinal axis clearly visible in a morph between the density maps of bare F-actin and rigor actomyosin-15 (Video S1). Consistently, the helical rise decreases 0.12 Å upon myosin-15 binding, with no substantial alteration of the helical twist (Figure S1C). The short *α*-helix in actin subunit i which forms contacts with D1647 shifts slightly towards the minus end to mediate this compression (Figure 4B). This is accommodated by rearrangements in the D-loop of subunit i + 2, which is also engaged by the myosin-15 HLH.

**Figure 4.**
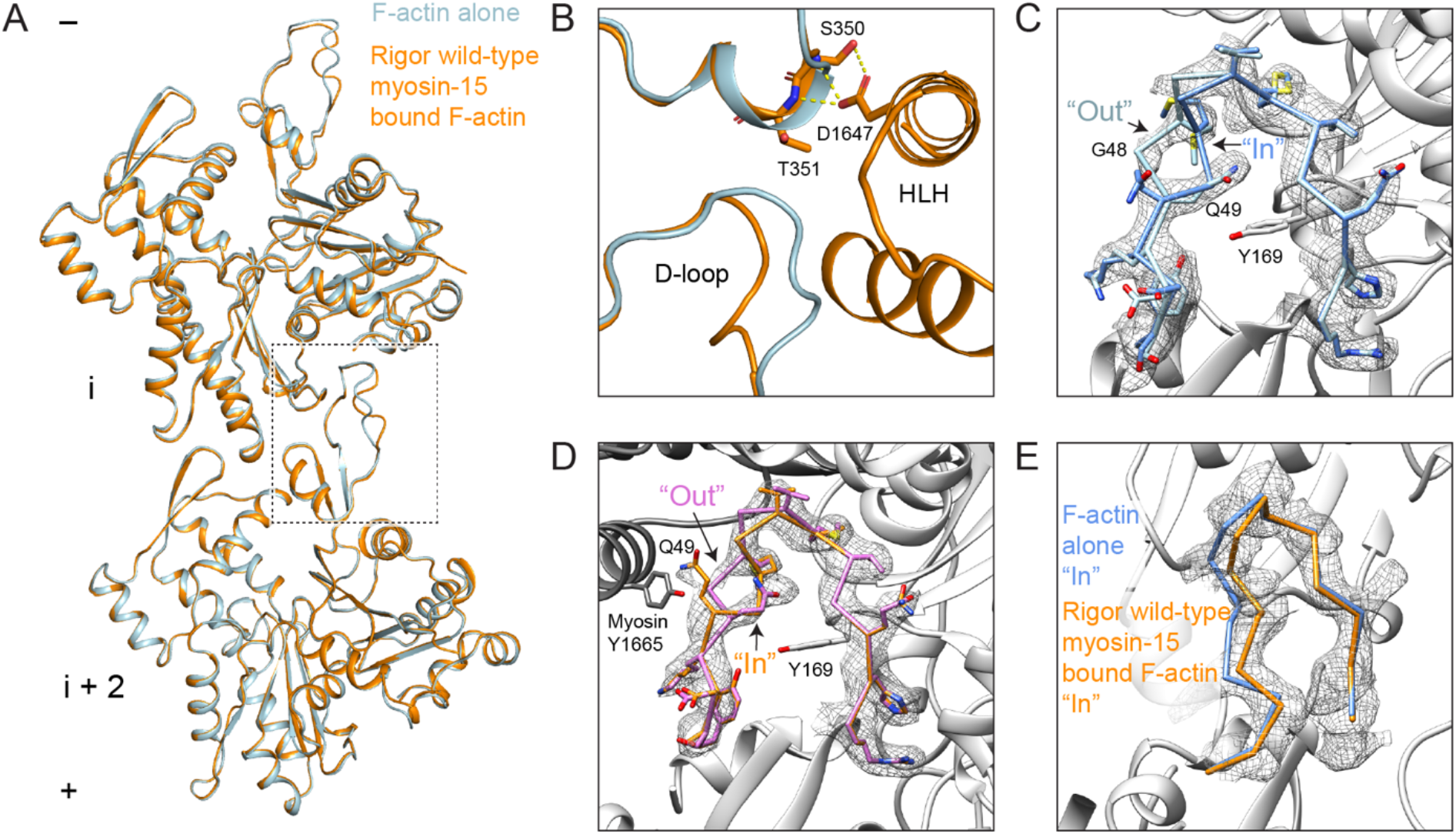
Myosin-15 binding remodels F-actin while maintaining D-loop flexibility. **(A)** Ribbon diagram of two actin subunits in the presence and absence of myosin-15, superimposed on subunit i. **(B)** Structural comparison at the interface formed by two longitudinally adjacent actin subunits and the myosin-15 HLH. Bare F-actin and rigor wild-type actomyosin-15 are colored in light blue and dark orange, respectively. **(C)** Segmented cryo-EM density of the bare F-actin D-loop rendered at 0.036 RMS, superimposed with stick models of the “Out” and “In” D-loop conformations colored in light blue and cornflower blue, respectively. **(D)** Segmented cryo-EM density of the rigor wild-type myosin-15 bound actin D-loop, superimposed with stick models of the “Out” and “In” D-loop conformations colored in dark orange and pink, respectively. **(E)** Backbone representation of the “In” conformation of the indicated models, superimposed on the segmented cryo-EM density the of myosin-15 bound F-actin D-loop.

Several recent high-resolution cryo-EM studies of F-actin have indicated that the D-loop is structurally dynamic within the context of the filament (Chou and Pollard, 2019; Das et al., 2020; Merino et al., 2018; Pospich et al., 2020). At a high threshold (0.036 RMS) our bare F-actin map features ambiguity surrounding residues G48 and Q49, despite clear density for all other residues in the D-loop (Figure 4C). However, at a lower threshold (0.025 RMS), the unprecedented resolution of our map reveals two continuous paths for the backbone, allowing two distinct conformations of the D-loop to be modeled: an “In” conformation where the main-chain of G48 flips in towards the center of the filament and Q49 faces the solvent, and an “Out” conformation where the main-chain of G48 is exposed and Q49 is buried in the center of the D-loop (Figures 4C, S5). Based on the respective strengths of the 2 density paths, the “Out” conformation is predominant in the bare F-actin structure, likely stabilized by an interaction between the side chain of Q49 of subunit i + 2 and the side chain of Y169 from the adjacent subunit i (Figure 4C). Upon rigor myosin-15 binding, the D-loop transitions to primarily being in the “In” state (Figure 4B,D), but displaced 2.7 Å outward from the filament axis relative to the bare F-actin conformation (Figure 4E), enabling an interaction between myosin HLH residue Y1665 and the flipped-out sidechain of actin D-loop residue Q49 (Figure 4D). Nevertheless, at a lower threshold (0.028 RMS), density resembling the “Out” conformation in bare actin is clearly visible (Figure 4D, Video S2), suggesting that rigor myosin-15 engages its binding site while maintaining F-actin’s intrinsic structural plasticity in the D-loop.

### The myosin-15 *jordan* deafness mutant locks the D-loop in the “In” conformation

To gain insight into structural mechanisms underlying the *jordan* mutant’s suppression of actin assembly and disruption of stereocilia growth (Moreland, 2021), we next compared the rigor structures of wild-type and *jordan* mutant actomyosin-15. The overall conformations of the motor domains are essentially indistinguishable, arguing the *jordan* mutation does not grossly compromise myosin-15’s structure (Figure S6A). The D1647G substitution abolishes the contacts formed between D1647 and actin residues S350 and T351 (Figure 5A) without disrupting the conformation and positioning of the HLH or discernibly perturbing any other contacts at the myosin-15-actin interface (Figure S6B-E). We next examined the structures for global differences in actin conformation, hypothesizing these could underlie the differential effects upon actin polymerization. Morphing the density maps reveals minimal changes, and the refined helical parameters of the reconstructions are nearly identical (Figure S1C and Video S3), indicating the *jordan* mutation does not substantially impact the global structural rearrangements in F-actin evoked by myosin-15. Consistently, the short *α*-helix comprising S350 and T351 of subunit i still undergoes a shift indistinguishable from that observed in the wild-type reconstruction (Figure 5B), indicating this conformational transition is not solely evoked by contacts formed with D1647 and is instead likely an allosteric effect driven by HLH engagement. Collectively, these data suggest that the *jordan* mutation also does not grossly disrupt the actin-myosin-15 interface or myosin-15’s conformational regulation of F-actin.

**Figure 5.**
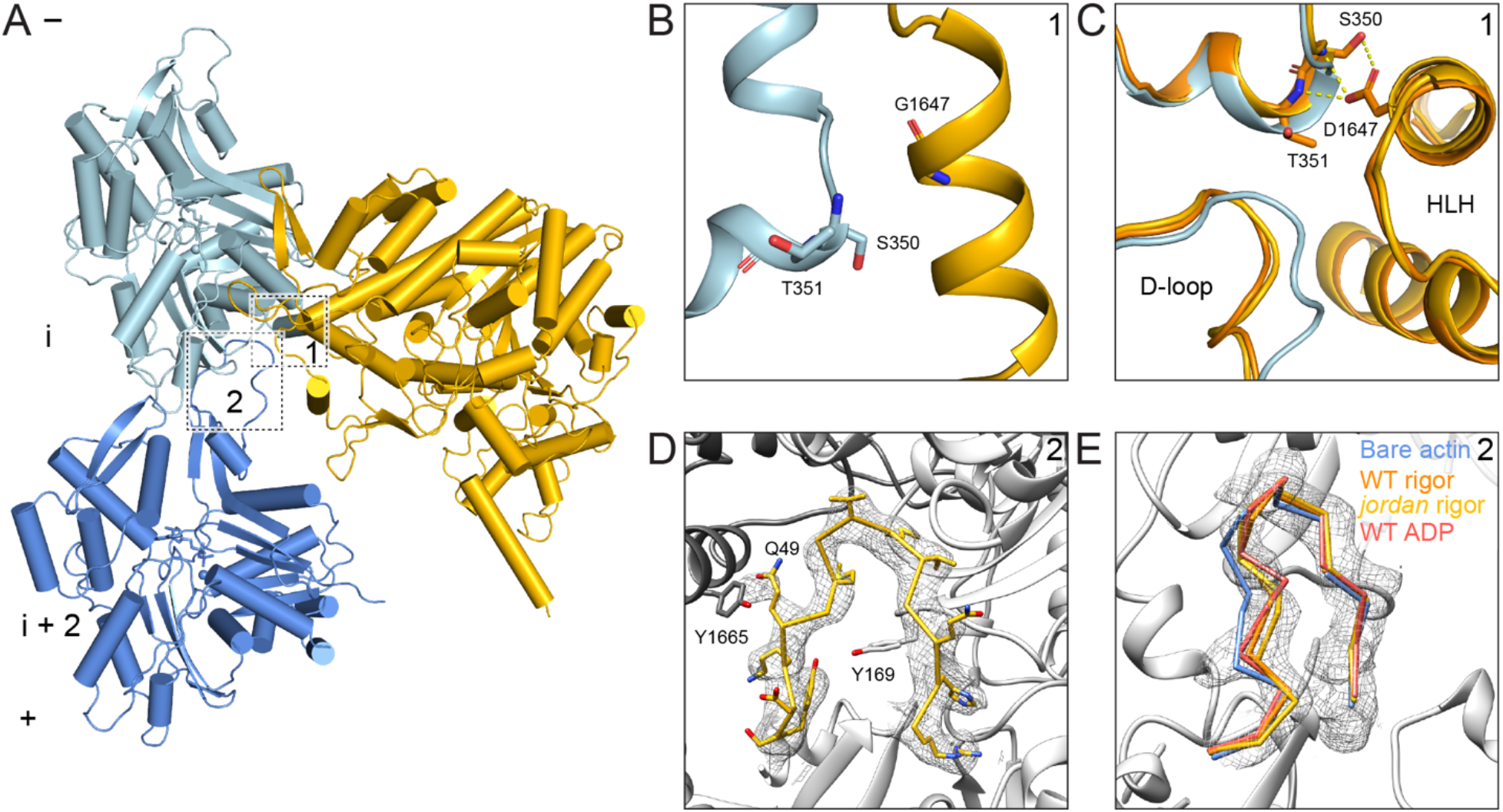
Binding by rigor *jordan* mutant myosin-15 locks the actin D-loop. **(A)** Atomic model of the interface between two actin subunits and rigor *jordan* myosin-15. Actin subunits i, i+2, and myosin-15 are colored in light blue, cornflower blue, and light orange, respectively. Numbered boxes correspond to detail views in subsequent panels. (**B**) Actomyosin interface at the site of the p.D1647G mutation. **(C)** Structural comparison of bare F-actin (light blue), rigor wild-type (dark orange), and *jordan* mutant (gold) myosin-15 bound F-actin at the interface formed by two longitudinally adjacent actin subunits and the myosin HLH. **(D)** Segmented cryo-EM density and atomic model of the rigor *jordan* mutant actomyosin-15 D-loop. **(E)** Backbone diagram of the three “In” D-loop conformation models superimposed on segmented cryo-EM density of the rigor *jordan* mutant actomyosin-15 D-loop.

We thus turned to an alternative hypothesis, that the *jordan* mutant could disrupt actin structural plasticity, rather than global actin conformation. Despite the lower resolution of *jordan* mutant versus the wild-type actomyosin-15 reconstruction, the D-loop is clearly resolved, and the backbone and sidechain of each residue can be fully fit into the density, adopting an “In” conformation indistinguishable from the wild-type reconstruction (Figure 5C,D). Density for the “Out” conformation, on the other hand, is completely absent, demonstrating that the *jordan* mutant locks the D-loop in the “In” conformation and eliminates its flexibility (Video S4). Thus, the *jordan* mutation appears to unexpectedly restrict the structural plasticity of the D-loop, presumably through a long-range allosteric effect, rather than eliciting structural rearrangements immediately adjacent to the site of the p.D1647 lesion. While we cannot rule out the possibility that the *jordan* mutation elicits additional perturbations of myosin-15 or F-actin structure during other steps of the myosin mechanochemical cycle, wild-type myosin-15 most potently stimulates F-actin assembly in the absence of nucleotide, arguing the *jordan* mutant’s defect likely occurs in a strongly-bound state (Moreland, 2021). As restricted D-loop flexibility is the only distinguishable difference between the two rigor structures, we propose that this dysregulation of actin structural plasticity contributes to inhibition of actin assembly by the *jordan* mutant (Moreland, 2021). Our structures furthermore support a key role for actin residue G48 in conferring D-loop flexibility mediating F-actin assembly. In *γ*-actin, the p.G48R substitution (Figure S3B) has been documented to cause progressive hearing loss in human patients (Miyagawa et al., 2015), phenocopying the *jordan* mutation. While D-loop flexibility at this position is likely a general feature of mammalian actin isoforms (Graceffa and Dominguez, 2003; Otterbein et al., 2001; von der Ecken et al., 2015), our data suggest that it is specifically required for efficient actin polymerization in stereocilia.

### The Mg-ADP state of myosin-15 stabilizes the D-loop to restrict actin nucleation

The ability of myosin-15 to nucleate F-actin is potently reduced by the presence of ATP (Moreland, 2021), leading us to hypothesize that other states in the motor’s mechanochemical cycle could have distinct effects on actin polymerization by differentially regulating F-actin conformation, as we have previously reported for myosin-6 (Gurel et al., 2017). We next undertook a detailed comparison of the rigor and Mg-ADP-bound actomyosin-15 wild-type structures. The overall conformation of the myosin-15 motor domain is highly similar in both reconstructions, with a global RMSD of 0.462 Å for 493 aligned C*_α_* atoms (Figure 6A), indicating an absence of major subdomain rearrangements during ADP release (Figure S7A). Clear density corresponding to ADP is present in the Mg-ADP state cryo-EM map (Figure 6A), arguing that the observed lack of rearrangements is not due to insufficient nucleotide binding by the motor. Both myosin-1 and myosin-6 feature a minor lever-arm swing accompanying ADP release, which has been postulated by us and others to mediate the known force-sensitivity of this step in their mechanochemical cycles (Gurel et al., 2017; Mentes et al., 2018; Nyitrai and Geeves, 2004). As the myosin-15 lever arm does not undergo a substantial transition (Figures 6A, S7A), we speculate that ADP release by myosin-15 is unlikely to be strongly mechanically-gated.

**Figure 6.**
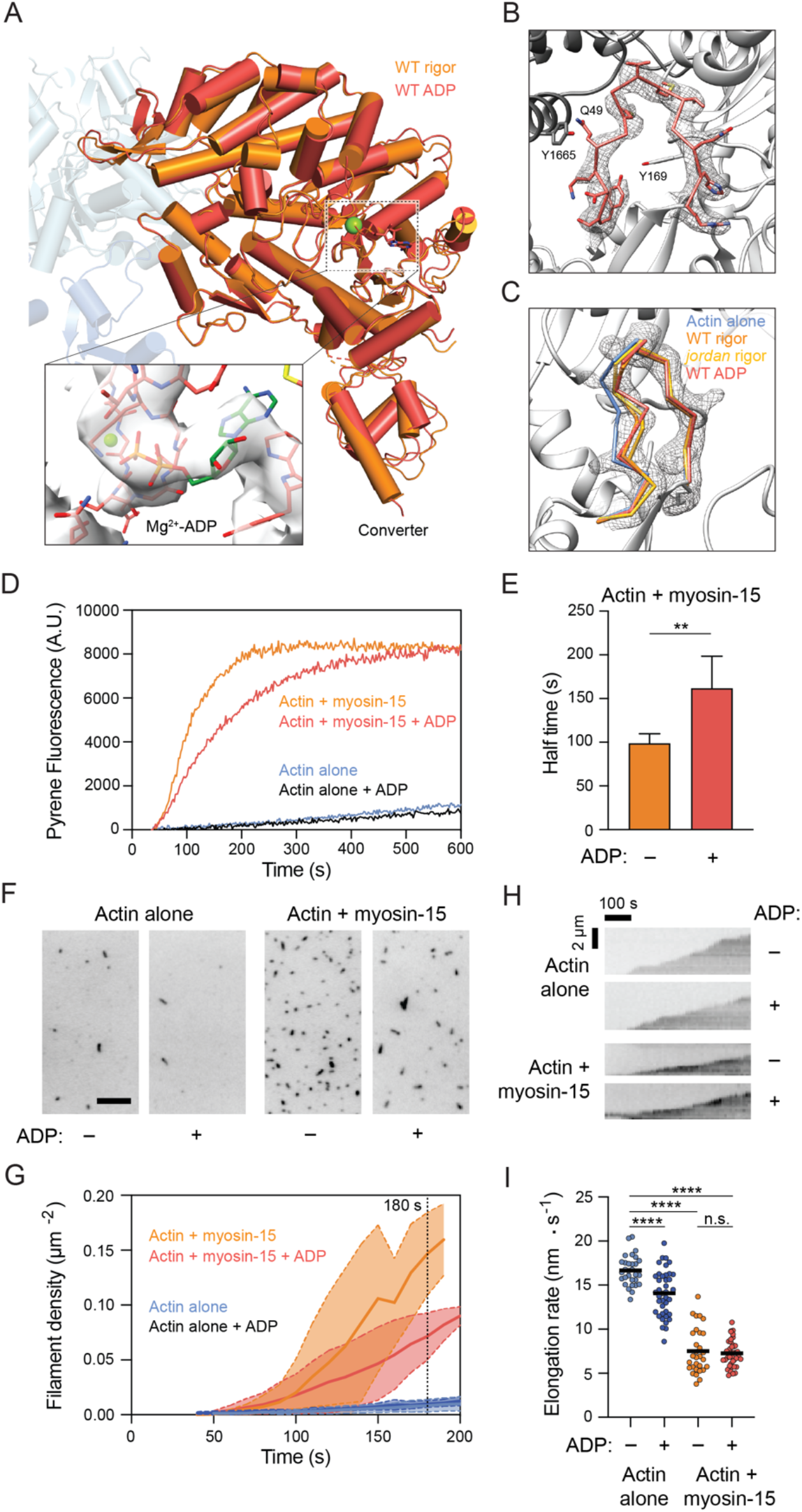
ADP bound myosin-15 locks the D-loop, blunting F-actin stimulation. **(A)** Superposition of rigor and ADP wild-type actomyosin-15 atomic models. Actins from the rigor atomic model are shown in shades of transparent blue. Inset: Cryo-EM density in the nucleotide-binding pocket of ADP wild-type myosin-15. **(B)** Cryo-EM map and corresponding atomic model of actin’s D-loop in ADP wild-type actomyosin-15. **(C)** Backbone positioning of “In” D-loop conformation in the indicated models, superimposed on the segmented cryo-EM map of the ADP wild-type actomyosin-15 D-loop. **(D)** Representative pyrene actin polymerization assays in the presence and absence of myosin-15 and ADP. Actin, 2 μM; myosin-15, 1 μM; Mg-ADP, 100 μM. **(E)** Quantification of the time to half-maximal pyrene signal saturation in the presence of myosin-15 with or without ADP. Error bars represent S.D. N = 5 from 3 independent protein purifications; **p = 0.008, Mann Whitney U-test. (F) Snapshots after 180 s of actin polymerization from TIRF movies recorded in the presence and absence of myosin-15 and ADP. Scale bar, 10 μm. **(G)** Quantification of F-actin density, indicative of nucleation rate, from TIRF movies. Solid lines and dashed lines / shading represent mean ± S.D. at each time point; N = 3. **(H)** Representative kymographs of elongating individual actin filaments from TIRF movies. **(I)** Quantification of filament elongation rates from TIRF movies. N = 3 independent replicates. Bars represent mean; ****p < 0.0001, one-way ANOVA.

We next compared the conformation of F-actin between the rigor and Mg-ADP structures. Consistent with the lack of conformational rearrangements in the motor domain, ADP release by myosin-15 evokes minimal global conformational changes in F-actin (Video S5); correspondingly, the helical rise of the two reconstructions is nearly identical (Figure S1C). Density for the D-loop is clearly resolved in the “In” conformation, positioned as we observed in both the wild-type and *jordan* mutant rigor reconstructions. However, like the *jordan* mutant actomyosin-15 reconstruction, density for the “Out” conformation is completely absent (Figure 6B-C, Video S6). As rigor *jordan* myosin-15 suppresses both D-loop dynamics and actin polymerization, we hypothesized ADP binding would correspondingly reduce the actin-polymerization stimulating activity of wild-type myosin-15 by also stabilizing the D-loop in the “In” conformation.

Consistent with this prediction, we observe a slower rate of actin assembly when saturating Mg-ADP is included along with myosin-15 in pyrene actin polymerization assays versus nucleotide-free conditions (Figure 6D), with an approximate doubling of the half-time to reach steady-state (Figure 6E). Inclusion of Mg-ADP did not substantially affect the assembly rate of actin alone (Figure 6D), suggesting ADP downregulates the polymerization-stimulating activity of myosin-15 rather than perturbing F-actin formation. We next sought to determine if ADP binding by myosin-15 impacted the nucleation or elongation phases of actin polymerization by directly visualizing filament dynamics with Total Internal Reflection Fluorescence (TIRF) microscopy (Figure 6F). Relative to actin alone, we observed a striking increase in the density of filaments per unit area in the presence of myosin-15, which was blunted by the inclusion of saturating Mg-ADP (Figure 6F,G), suggesting a specific effect on actin nucleation. When the growth of individual filament plus ends was monitored (Figure 6H), both nucleotide-free and Mg-ADP myosin-15 significantly slowed elongation to ∼7.5 nm / s, relative to ∼16 nm / s for actin alone (Figure 6I). This rate was indistinguishable from that observed in the presence of the rigor *jordan* mutant in our companion study (Moreland, 2021). As a further control, the addition of ADP only modestly affected the elongation rate of actin alone (Figure 6I). Myosin-15’s opposing effects of stimulating nucleation while reducing elongation strongly suggest that the overall increased F-actin assembly rate is dominated by myosin-15’s effect on stimulating nucleation, as previously reported for muscle myosin S1 (Fievez et al., 1997; Lheureux et al., 1993), in a manner which is specifically regulated by myosin-15 nucleotide state.

## Discussion

Here, we establish structural mechanisms underpinning myosin-15’s function in constructing the actin core of stereocilia, a process that is critical for hearing. Our actomyosin-15 cryo-EM structures, the first myosin-15 motor domain structures, to our knowledge, of any kind, assign the likely mechanistic deficiencies associated with many DFNB3 deafness mutations, as well as provide a framework for interpreting novel clinical variants (Figure 2). Many mutations are intuitively anticipated to impact the motor’s mechanochemical cycle. However, our structure also highlights surface substitutions which are not readily interpreted through this lens, suggesting additional functions of the myosin-15 motor domain important for hearing remain to be identified, such as serving as a protein-protein interaction hub. Our studies additionally reveal how myosin-15 modulates actin’s structural landscape to enhance F-actin nucleation in a manner controlled by the motor’s nucleotide state. We speculate this activity could be harnessed to provide fine-tuned control of F-actin assembly at stereocilia tips, facilitating the precise control of stereocilia height required for sound detection.

Our cryo-EM reconstructions and associated functional data reveal subtle structural plasticity in the actin D-loop which mediates regulated F-actin assembly by myosin-15. While substantial D-loop rearrangements accompanying the soluble G-actin to polymerized F-actin conformational transition are well-established (Dominguez and Holmes, 2011), the extent and functional role of D-loop rearrangements within F-actin remain controversial. Substantially different “open” and “closed” conformations have been reported in cryo-EM reconstructions of F-actin in the presence of stabilizing drugs and actin nucleotide-state analogs (Merino et al., 2018; Pospich et al., 2020) leading to speculation that an open-to-closed transition is associated with nucleotide hydrolysis and phosphate release concomitant with F-actin polymerization. However, others have reported minimal rearrangements when comparing highly similar conditions, with the D-loop constitutively adopting a closed conformation in F-actin (Chou and Pollard, 2019; Das et al., 2020). The data we present here reveal the closed D-loop conformation in ADP F-actin to be a mixture of “In” and “Out” states whose occupancy can be manipulated by myosin-15 engagement. Wild-type rigor myosin-15 binding retains both the “In” and “Out” D-loop conformations and strongly stimulates actin polymerization, while ADP-bound wild-type myosin-15 and rigor *jordan* mutant-15 both lock the D-loop in the “In” state, with corresponding moderate and striking reductions, respectively, in F-actin-polymerization stimulation activity (Figure 7). Collectively, these data suggest that the co-existence of the “In” and “Out” D-loop conformations promotes polymerization relative to locking the D-loop in the “In” conformation, a model which is consistent with a recent report demonstrating that constraining D-loop flexibility with short-distance crosslinkers was refractory to actin polymerization and stimulated F-actin disassembly (Das et al., 2020). Previous studies have further shown that muscle myosin S1 can rescue polymerization of actin featuring a proteolytically disrupted D-loop (Wawro et al., 2005), consistent with a role for myosin in modulating D-loop dynamics during actin polymerization.

**Figure 7.**
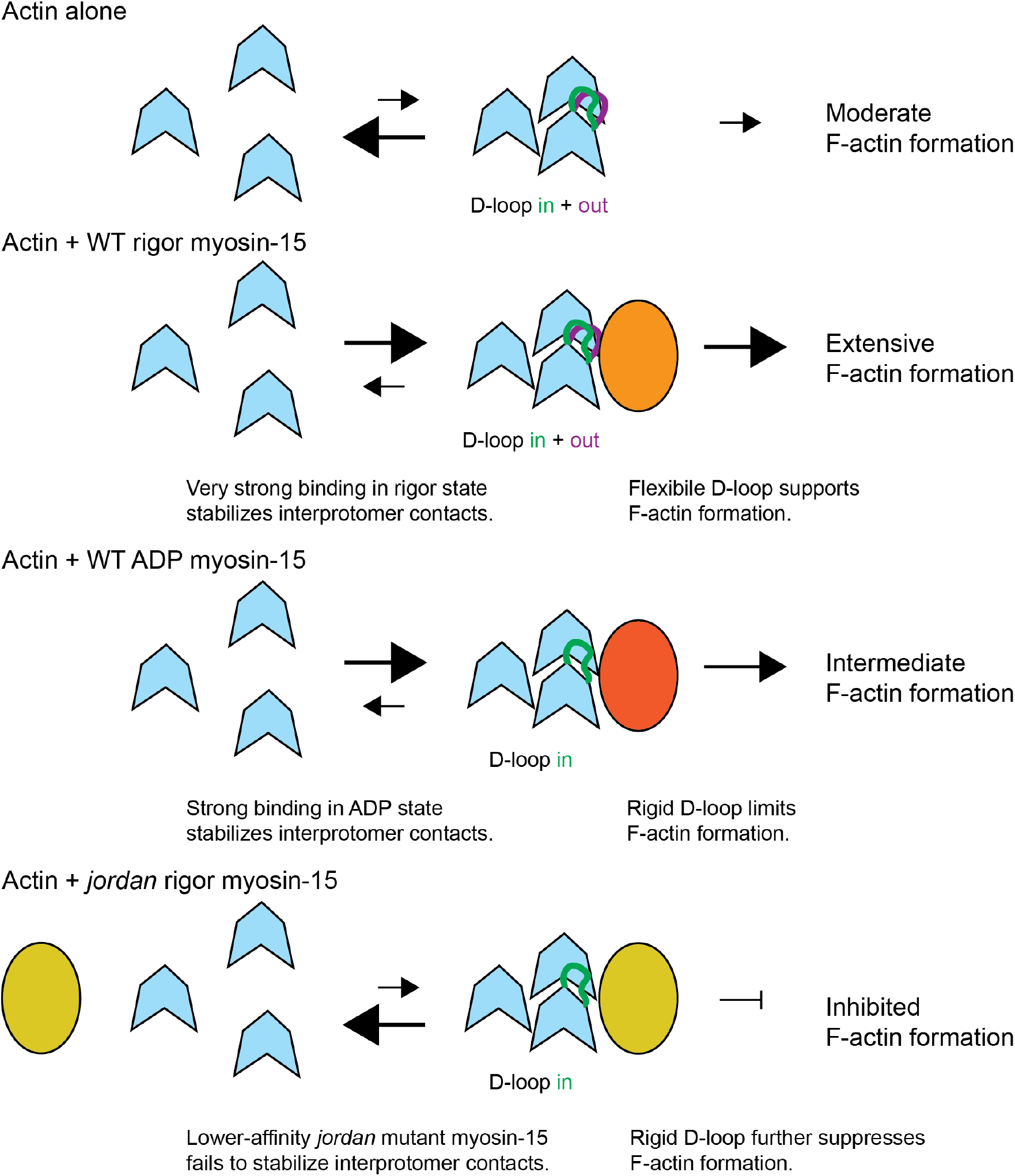
Schematic model for regulation of actin polymerization by myosin-15. We propose myosin-15 enhances F-actin formation by stimulating nucleation, both by stabilizing actin-actin contacts and modulating D-loop structural plasticity. Myosin-15 nucleotide state facilitates tuning polymerization-stimulation activity, which could be harnessed for stereocilium height control. The *jordan* mutation disrupts F-actin regulation, thereby inhibiting polymerization.

Nevertheless, additional mechanisms are likely required to explain wild-type myosin-15’s stimulation of actin polymerization and its striking disruption by the *jordan* deafness mutation. While actin polymerization by muscle myosin S1 has been subjected to extensive biochemical scrutiny, the exact mechanism remains controversial. Chaussepied and colleagues have suggested that each S1 engages a single G-actin, which then stimulates polymerization through a standard nucleation-elongation mechanism (Lheureux et al., 1993). Carlier and co-workers, on the other hand, have suggested that each S1 engages two G-actins, forming trimers with an F-actin-like actin-actin longitudinal bond which coalesce into filaments through a mechanism distinct from standard polymerization (Fievez et al., 1997). While our data do not directly discriminate between these mechanisms, our observation that the *jordan* mutation is localized to the HLH interface, which bridges the contact between two longitudinally-adjacent protomers, supports a model in which high-affinity myosin-15 binding promotes longitudinal interactions to stimulate polymerization (Figure 7). The modestly lower affinity of ADP-myosin 15 and strikingly lower affinity of the *jordan* mutant for F-actin, relative to the rigor wild-type (Moreland, 2021), also likely makes a major contribution to determining their potency in stimulating F-actin formation. This defect occurring at the level of nuclei formation, rather than elongation, is consistent with our observation that F-actin elongation rates are indistinguishable in the presence of ADP-bound or rigor wild-type myosin-15 (Figure 6H and I), as well as the rigor *jordan* mutant (Moreland, 2021), all of which slow actin polymerization relative to G-actin alone. Regardless, diminished affinity of the *jordan* mutant cannot be the sole explanation for its polymerization defect, as the presence of this myosin, which nevertheless binds F-actin with measurable affinity in an almost identical pose to the wild-type, actually initially suppresses the rate of F-actin formation relative to G-actin alone (Moreland, 2021). The most parsimonious explanation for this effect is through the suppression of D-loop flexibility as highlighted above, emphasizing the contributions of both myosin-15 bridging actin monomers to stimulate nucleus formation, and the maintenance of actin conformational plasticity to the accelerated actin polymerization we observe (Figure 7).

Our studies define a biophysical and structural framework for tunable stimulation of actin polymerization by myosin-15 which could be modulated by the local biochemical environment at the tips of stereocilia, the major sites of actin polymerization. We speculate that hair cells harness the differential efficiency of actin nucleation by myosin-15 in distinct nucleotide states as a regulatable mechanistic component of stereocilia height control. It is feasible that the ADP-bound state, which substantially accelerates actin polymerization, albeit less than the rigor state, is sufficient for supporting myosin-15’s contribution to actin polymerization in the tip compartment. Within this framework, control over myosin-15’s actin nucleation could be exerted by the availability of ATP versus ADP (Moreland, 2021), which has previously been reported to be dynamically maintained by creatine kinase in stereocilia during mechanotransduction and adaptation (Shin et al., 2007) in a manner which may vary in different species and hair cell subtypes (Krey and Barr-Gillespie, 2019). Our structural data furthermore suggest that ADP release is unlikely to be mechanically-gated in myosin-15, which could facilitate the motor populating a nucleotide-free state in the cell in order to boost its nucleation activity in certain ATP / ADP concentration regimes. In future studies, it will be important to determine the dynamic availability of ATP and ADP in the tip compartment of developing stereocilia, and how myosin-15’s direct effects on actin polymerization are coordinated with the WHRN-EPS8-GPSM2-GNAI3 elongation network that it transports. Other environmental conditions known to impact stereocilium integrity, such as oxidative stress (Wagner and Shin, 2019), could also potentially impact stimulated actin polymerization by myosin-15, both by dysregulation of nucleotide availability and by direct modification of the myosin-15 protein, disrupting the actin-myosin interface in a manner conceptually analogous to the *jordan* mutation. Finally, our observation that the *jordan* mutation occurs at a conserved position suggests that other unconventional myosins could also feasibly stimulate actin polymerization in stereocilia, as well as in diverse physiological contexts beyond hair cells. The mechanistic framework we establish here will broadly facilitate investigating the functional role of unconventional myosins in actin assembly dynamics.

## Acknowledgements

We gratefully acknowledge Yasuharu Takagi and James Sellers (NHLBI DIR) for assistance with myosin biochemistry, Rabia Faridi (NIDCD DIR) for a curated list of DFNB3 mutations, and Peter Barr-Gillespie for critical feedback. We also thank Johanna Sotiris and Mark Ebrahim from the Rockefeller University Cryo-EM Resource Center (CEMRC) for their assistance with data collection. R.G. was supported by an H. Li Memorial Fellowship and P.G. by a Rockefeller Women in Science Fellowship. This research was supported (in part) by the Intramural Research Program of the NIH, NIDCD DC000039 to Thomas B. Friedman. The work was additionally funded by grants from the Medical Research Council to M.R.B. (MC_UP_1503/2), the Pew Charitable Trusts and Irma T. Hirschl / Monique Weill-Caulier Trust to G.M.A., as well as National Institutes of Health grants to J.E.B (R01DC018827) and G.M.A. (DP5OD017885 and R01GM141044). The content of this manuscript is solely the responsibility of the authors and does not necessarily represent the official views of the National Institutes of Health.

## Author Contributions

**R.G.:** Conceptualization, Methodology, Investigation, Formal analysis, Visualization, Writing - Original draft. **F.J.:** Investigation, Formal analysis. **Z.G.M.:** Investigation, Formal analysis. **M.J.R.:** Methodology, Formal analysis. **S.E.R.:** Investigation, Formal analysis. **P.G.** Methodology, Investigation. **A.S.:** Methodology, Investigation. **M.R.B.:** Resources. **J.E.B.:** Conceptualization, Visualization, Project administration, Funding acquisition, Writing - Original draft, Supervision: **G.M.A.** Conceptualization, Formal analysis, Visualization, Project administration, Funding acquisition, Writing - Original draft, Supervision. **All authors:** Writing – Review & editing.

## Data Availability Statement

Cryo-EM maps and corresponding atomic models have been deposited in the EMDB and PDB databases. Accession codes are as follows: rigor wild-type myosin-15 bound F-actin: EMDB: EMD-24322, PDB: 7R91; rigor *jordan* myosin-15 bound F-actin: EMDB: EMD-24400, PDB: 7RB9; ADP wild-type myosin-15 bound F-actin: EMDB: EMD-24399, PDB: 7RB8; F-actin alone: EMDB: EMD-24321, PDB: 7R8V.

---

**Figure S1.**
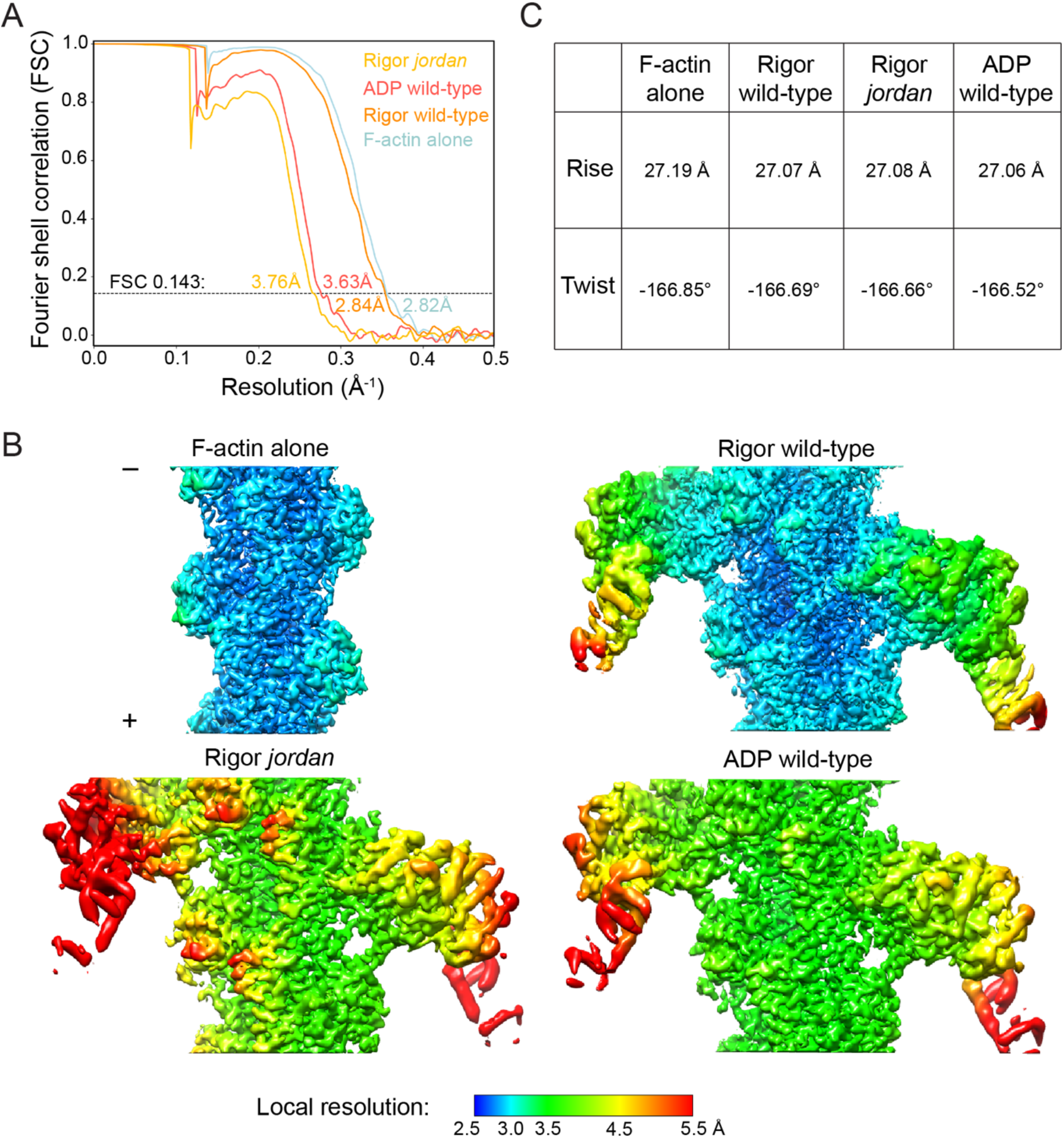
Resolution assessment of cryo-EM reconstructions and helical parameters. **(A)** Gold-standard Fourier shell correlation (FSC) curves for all 3D reconstructions presented in this study. Overall resolution is estimated by the FSC 0.143 criterion. **(B)** Local resolution estimation of the indicated reconstructions. **(C)** Helical parameters of the 3D reconstructions.

**Figure S2.**
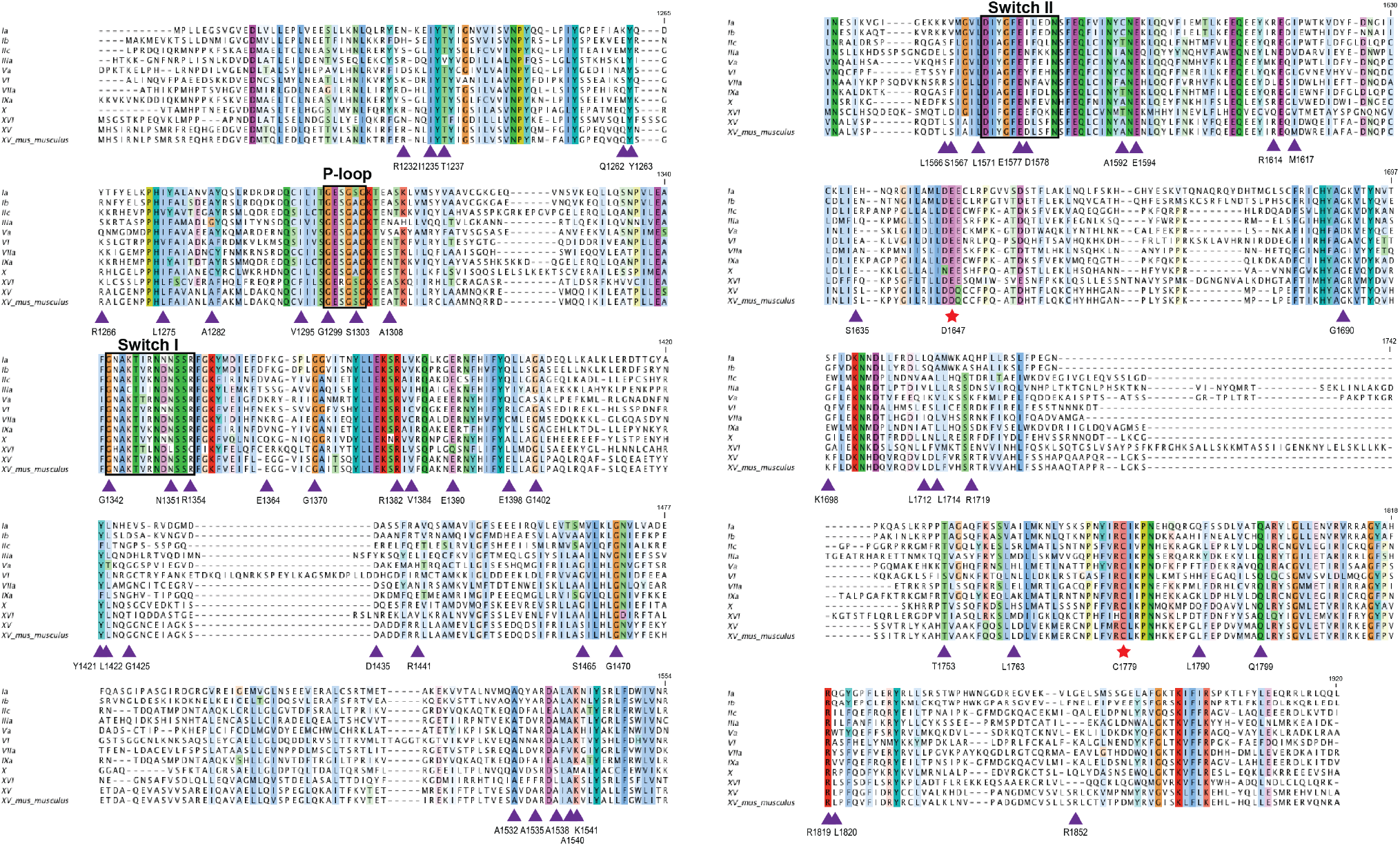
Sequence alignment of myosin motor domains. Aligned sequences correspond to: *H. sapiens* myosin-1a (NP_005370.1), 1b (NP_001155291.1), 2c (NP_079005.3), 3a (NP_059129.3), 5a (NP_000250.3), 6 (NP_004990.3), 7a (NP_000251.3), 9a (NP_002464.1), 10 (NP_036466.2), 16 (NP_001185879.1), 15 (NP_057323.3), and *M. musculus* 15 (NP_874357). The alignment is colored by sequence conservation. The sites of D1647 and C1779 are indicated by red stars, and sites of deafness mutations are indicated by purple triangles. Sequence alignment was performed using Clustal Omega and formatted with Jalview.

**Figure S3.**
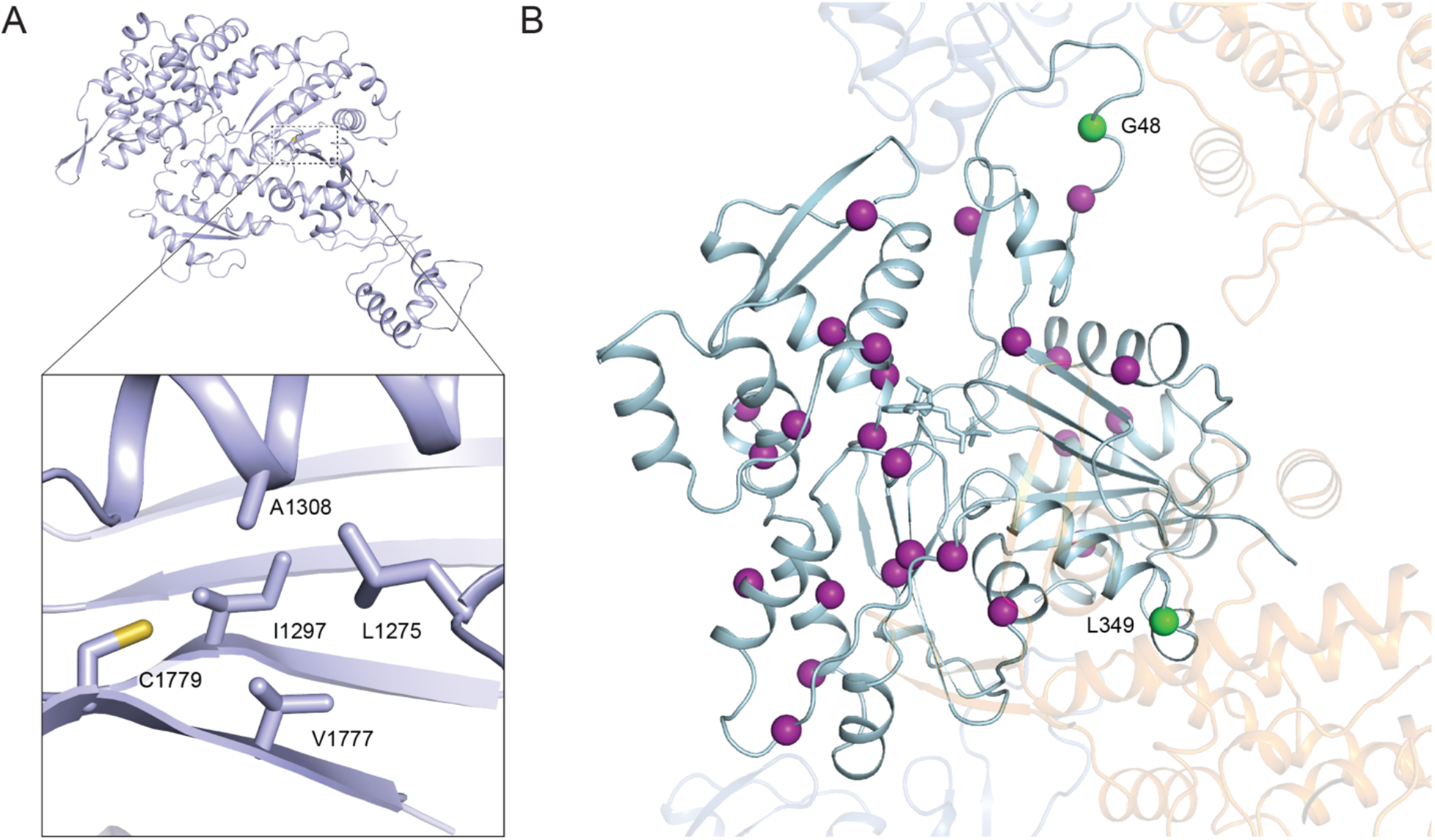
Additional structural analysis of deafness causing mutations. **(A)** The hydrophobic environment surrounding C1779, the site of the *shaker 2* mutation. Hydrophobic residues are shown as sticks. **(B)** Deafness causing mutations on *γ*-actin shown as magenta spheres. Two residues located at the actomyosin-15 interface are highlighted in green. Adjacent actin subunits and myosin-15 are shown in transparent blue and orange, respectively.

**Figure S4.**
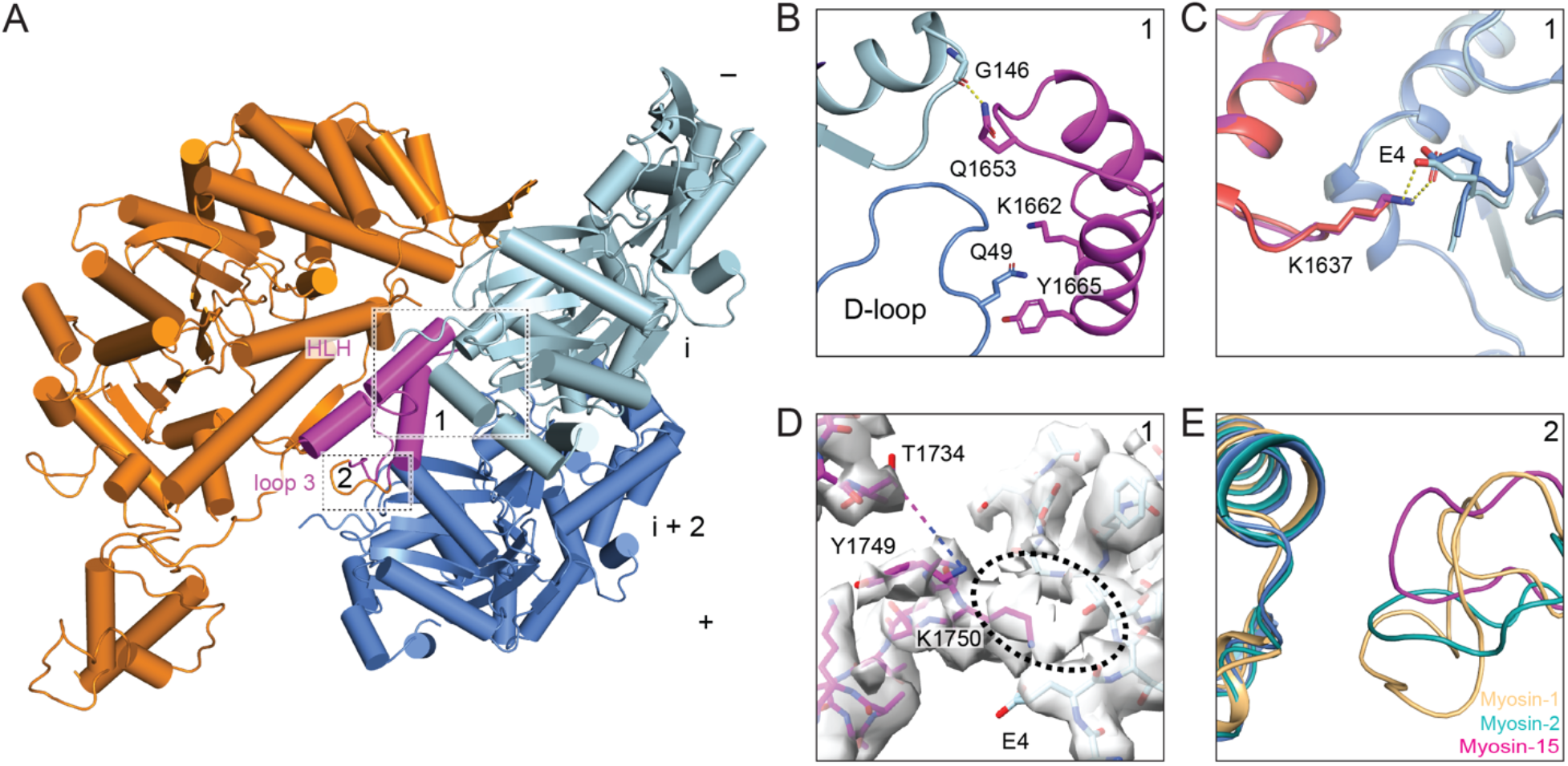
Additional interactions at the actomyosin-15 interface. **(A)** Atomic model of the interface between two actin subunits and rigor wild-type myosin-15 shown from an alternative perspective. Actin subunits i, i+2, and myosin-15 are colored in light blue, cornflower blue, and dark orange, respectively. The indicated actin interaction motifs are highlighted in magenta, and numbered boxes correspond to detail views in subsequent panels. **(B)** Electrostatic interactions between the HLH of rigor wild-type myosin-15 and F-actin. **(C)** Interaction between the activation loop of myosin-15 and the N terminus of actin subunit i. The HLH of the rigor and ADP wild-type myosin-15 are colored in magenta and deep salmon. The corresponding actin subunits are colored in light blue and cornflower blue. **(D)** The density map around loop 2 of rigor wild-type myosin-15. The models of myosin-15 and actin are colored in magenta and light blue. The disordered region of loop 2 between residues T1734 and Y1749 is represented by a dashed line. Extra density near the N terminus of actin is circled. **(E)** Structural comparison of the myosin loop 3-actin interface between rigor state myosin-15, myosin-1, and myosin-2. The structures are superimposed on the loop 3-interacting actin subunit. Loop 3 of myosin-15 is colored in magenta and the interacting actin is colored in cornflower blue, while loop 3 of myosin-1 and myosin-2 are colored in light orange and teal, respectively, as are their interacting actin subunits.

**Figure S5.**
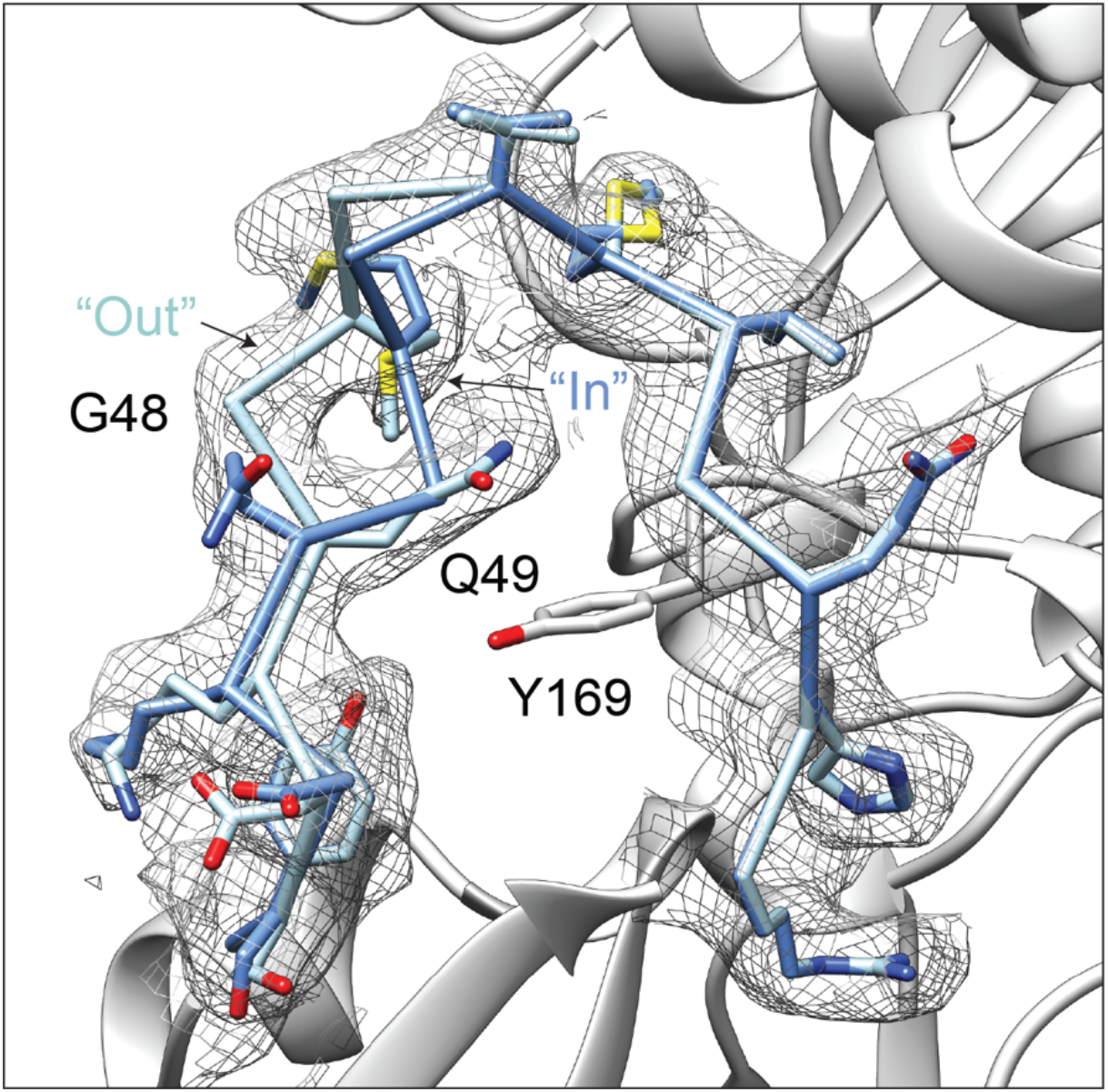
The D-loop of bare F-actin assumes two conformations. Magnified view of segmented cryo-EM density of the bare F-actin D-loop rendered at 0.025 RMS, superimposed with stick models of the “Out” and “In” D-loop conformations colored in light blue and cornflower blue, respectively.

**Figure S6.**
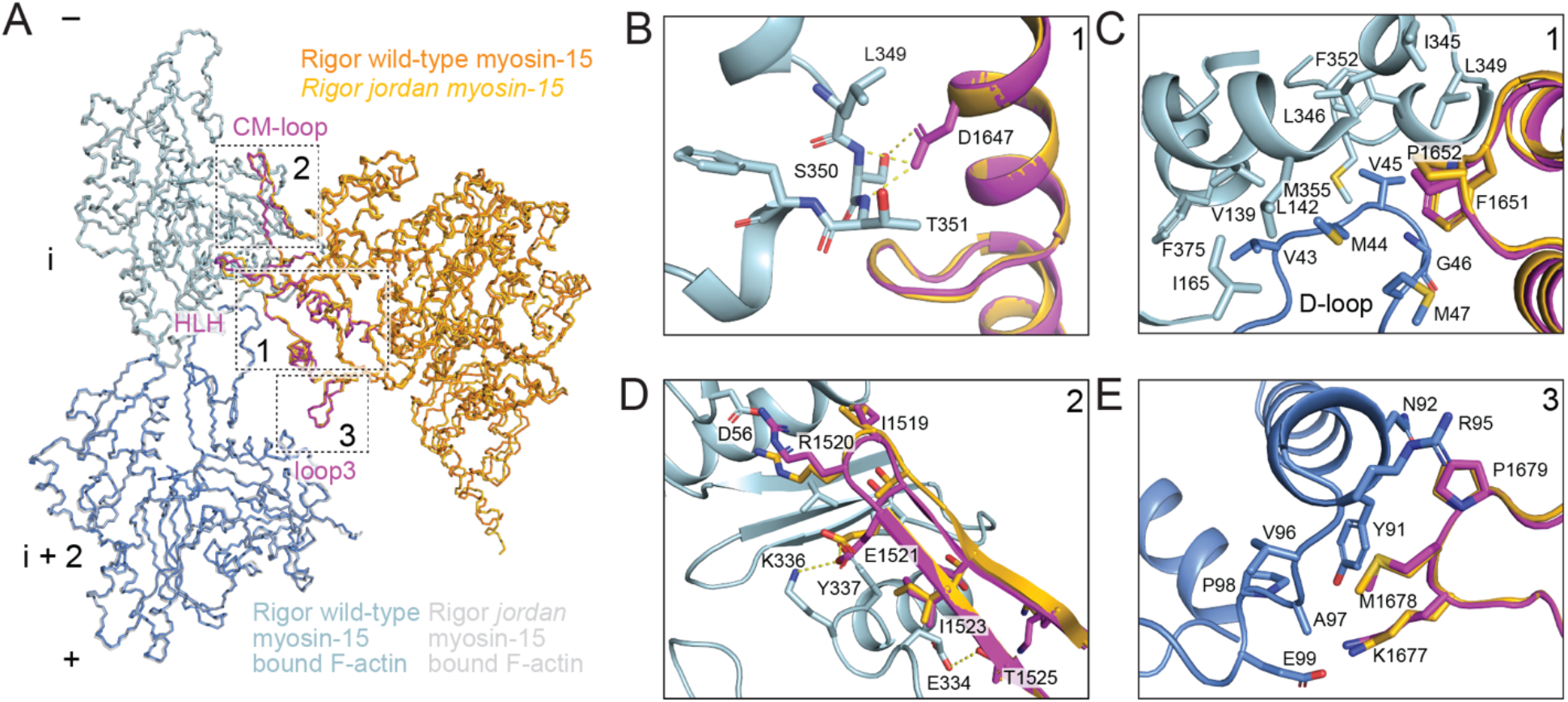
Structural comparison of rigor wild-type and *jordan* mutant actomyosin-15 at the actin-myosin interface. **(A)** Backbone representations of the rigor wild-type and *jordan* mutant actomyosin-15 models, superimposed on actin subunit i. Myosin-15’s actin interacting motifs are highlighted in magenta on the rigor wild-type model. Numbered boxes correspond to detail views in subsequent panels, which match those displayed in Figure 3. **(B)** Although the D1647G lesion disrupts interactions in the *jordan* mutant, the overall HLH conformation in this region is maintained. **(C-E)** All other actin interface interactions mediated by the HLH (**C**), CM loop (**D**), and loop 3 (**E**), are maintained in the *jordan* mutant.

**Figure S7.**
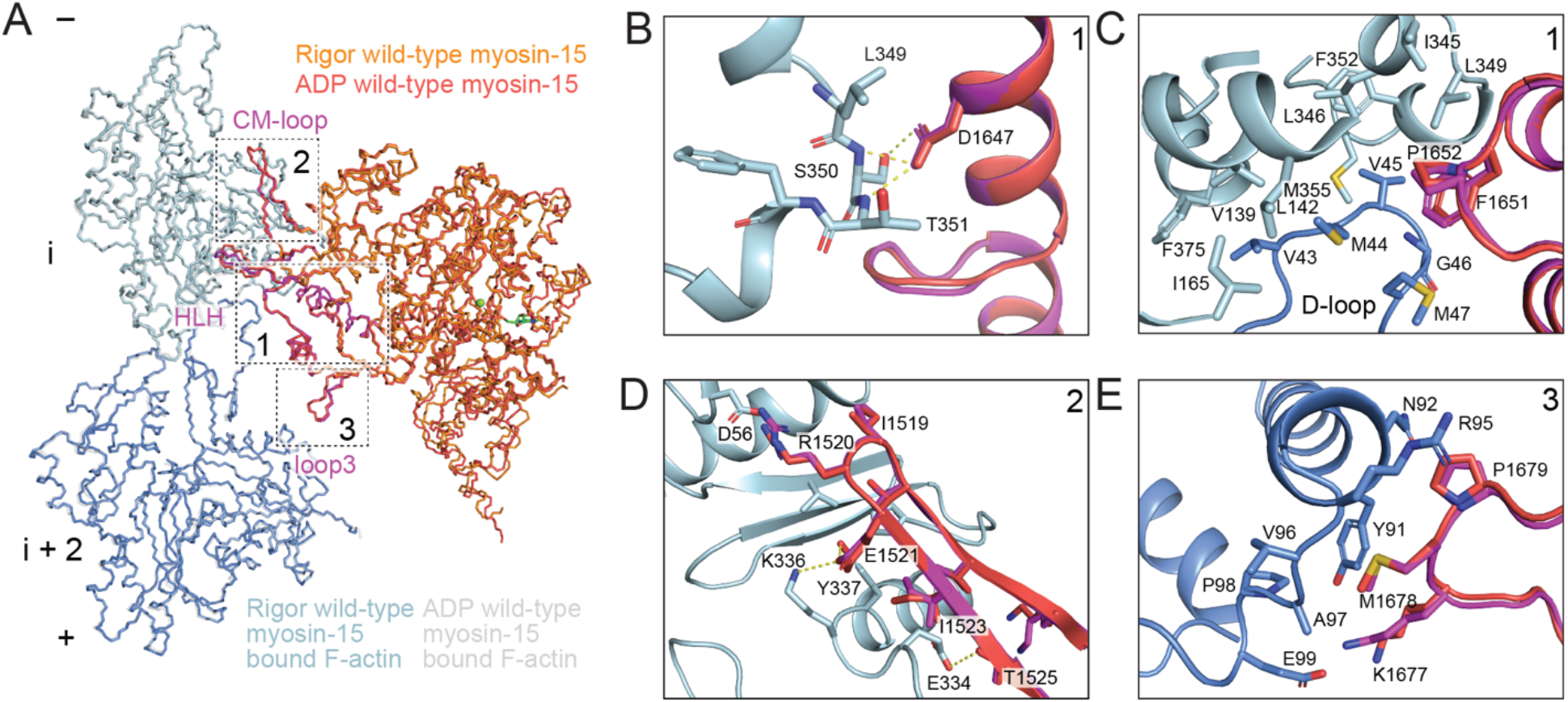
Structural comparison of rigor and ADP wild-type actomyosin-15 at the actin-myosin interface. **(A)** Backbone representations of the rigor and ADP wild-type actomyosin-15 models, superimposed on actin subunit i. Myosin-15’s actin interacting motifs are highlighted in magenta on the rigor wild-type model. Numbered boxes correspond to detail views in subsequent panels, which match those displayed in Figure 3. **(B-E)** All actin interface interactions mediated by HLH residues D1647 (**B**) and F1651/P1652 (**C**), as well as the CM loop (**D**) and loop 3 (**E**), are present in both the ADP and rigor states.

**Table S1.** Cryo-EM data collection, refinement, and validation statistics.

|  | **F-actin alone**<br>EMD-24321, PDB 7R8V | **Rigor WT myosin-15**<br>**bound F-actin**<br>EMD-24322, PDB 7R91 | **Rigor *jordan* myosin-15**<br>**bound F-actin**<br>EMD-24400, PDB 7RB9 | **ADP WT myosin-15**<br>**bound F-actin**<br>EMD-24399, PDB 7RB8 |
| --- | --- | --- | --- | --- |
| **Data collection and processing** |  |  |  |  |
| Microscope | Titan Krios | Titan Krios | Titan Krios | Titan Krios |
| Voltage (kV) | 300 | 300 | 300 | 300 |
| Detector | K2 Summit | K2 Summit | K2 Summit | K2 Summit |
| Magnification | 29,000 | 29,000 | 29,000 | 29,000 |
| Electron exposure (e<sup>-</sup> / Å<sup>2</sup>) | 60 | 60 | 60 | 60 |
| Exposure rate (e<sup>-</sup> / pixel / s) | 1.5 | 1.5 | 1.5 | 1.5 |
| Calibrated pixel size (Å) | 1.03 | 1.03 | 1.03 | 1.03 |
| Defocus range (µm) | -1.5 to -3.5 | -1.5 to -3.5 | -1.5 to -3.5 | -1.5 to -3.5 |
| Helical symmetry | C1 | C1 | C1 | C1 |
|  | 27.19 Å rise | 27.07 Å rise | 27.08 Å rise | 27.07 Å rise |
|  | -166.85° twist | -166.69° twist | -166.66° twist | -166.52° twist |
| Initial filament segments (no.) | 229,833 | 150,424 | 113,662 | 139,547 |
| Final filament segments (no.) | 228,334 | 142,635 | 91,340 | 125,232 |
| Map resolution (Å) | 2.82 | 2.83 | 3.76 | 3.63 |
| FSC threshold | 0.143 | 0.143 | 0.143 | 0.143 |
| **Refinement** |  |  |  |  |
| Initial model (PDB ID) | 6BNO | 6BNO | 6BNO | 6BNO |
| Model resolution (Å) | 3.0 | 3.0 | 3.9 | 3.8 |
| FSC threshold | 0.5 | 0.5 | 0.5 | 0.5 |
| Map sharpening B factor (Å<sup>2</sup>) | -44.79 | -50.05 | -86.65 | -84.62 |
| Model composition | 5 actin protomers | 3 actin protomers,<br>1 myosin-15 | 3 actin protomers,<br>1 myosin-15 | 3 actin protomers,<br>1 myosin-15 |
| Non-hydrogen atoms | 14845 | 14320 | 14193 | 14226 |
| Protein residues | 1855 | 1786 | 1786 | 1786 |
| Ligands | 5 Mg.ADP | 3 Mg.ADP | 3 Mg.ADP | 4 Mg.ADP |
| B factors (Å<sup>2</sup>) |  |  |  |  |
| Protein | 37.16 | 54.6 | 120.24 | 79.79 |
| Ligand | 27.66 | 29.25 | 77.38 | 70.69 |
| R.M.S. deviations |  |  |  |  |
| Bond lengths (Å) | 0.004 | 0.004 | 0.004 | 0.004 |
| Bond angles (°) | 0.97 | 0.978 | 0.979 | 1.018 |
| **Validation** |  |  |  |  |
| MolProbity score | 1.44 | 1.37 | 1.7 | 1.84 |
| Clash score | 5.56 | 5.01 | 9.24 | 8.73 |
| Poor rotamers (%) | 0 | 0.19 | 0 | 0.07 |
| Ramachandran plot |  |  |  |  |
| Favored (%) | 97.27 | 97.45 | 96.67 | 94.62 |
| Allowed (%) | 2.73 | 2.55 | 3.33 | 5.38 |
| Disallowed (%) | 0.00 | 0.00 | 0.00 | 0.00 |

## Supplementary Video Captions

**Video S1: Morph of segmented actin density between F-actin alone and rigor wild-type myosin-15 bound F-actin.** Maps were lowpass filtered to 5 Å.

**Video S2: Detail view of D-loop in morph between F-actin alone and rigor wild-type myosin-15 bound F-actin.** Maps are identical to those presented in Video S1.

**Video S3: Morph of segmented actin density between rigor wild-type and *jordan* myosin-15 bound F-actin.** Maps were lowpass filtered to 5 Å.

**Video S4: Detail view of D-loop in morph between rigor wild-type and *jordan* myosin-15 bound F-actin**. Maps are identical to those presented in Video S3.

**Video S5: Morph of segmented actin density between rigor and ADP wild-type myosin-15 bound F actin.** Maps were lowpass filtered to 5 Å.

**Video S6: Detail view of D-loop in morph between rigor and ADP wild-type myosin-15 bound F-actin.** Maps are identical to those presented in Video S5.

## Methods

### Expression and purification of myosin-15

Baculoviruses encoding either the wild-type, or *jordan* variant of the mouse myosin-15 motor domain (NP_874357.2, aa. 1 - 743) truncated after the 2^nd^ IQ domain, and including a C-terminal EGFP and FLAG moiety, were produced as described (Moreland, 2021). Briefly, *Sf*9 cells were seeded at 2 x 10^6^ cells / mL in ESF-921 (Expression Systems), and infected simultaneously with myosin-15 baculovirus at a multiplicity of infection (MOI) of 5. Additional dual-promoter baculoviruses expressing bovine smooth muscle essential (MYL6) and chicken regulatory (MYL12B) light chains (MOI = 5), as well as mouse UNC45B and HSP90AA1 (MOI = 5) were included (Bird et al., 2014; Pato et al., 1996). Cells were harvested 48 - 72 hours post-infection and flash frozen in liquid nitrogen.

Myosin-15 motor domains were purified as described (Moreland, 2021). Briefly, cells were homogenized in Extraction Buffer: 10 mM MOPS, 500 mM NaCl, 1 mM EGTA, 10 mM MgCl_2_, 2 mM ATP, 0.2 mM PMSF, 0.1 mM DTT, 1 mM NaN_3_, 2 μg·mL^−1^ leupeptin, 1 x protease inhibitor cocktail (Halt EDTA-free; Thermo Scientific), pH 7.2. Lysates were clarified at 48,000 x *g* for 30 minutes and incubated with FLAG M2 affinity resin (Sigma-Aldrich) for 3 hours at 4 °C. The FLAG resin was then washed in High-Salt Buffer (10 mM MOPS, 500 mM NaCl, 1 mM EGTA, 5 mM MgCl_2_, 1 mM ATP, 0.1 mM PMSF, 0.1 mM DTT, 1 mM NaN_3_, 2 μg·mL^−1^ leupeptin, pH 7.2) followed by Low-Salt Buffer (10 mM MOPS, 60 mM NaCl, 1 mM EGTA, 0.1 mM PMSF, 0.1 mM DTT, 1 mM NaN_3_, 2 μg·mL^−1^ leupeptin, pH 7.0). Myosin-15 protein was subsequently eluted with 0.2 mg·mL^−1^ 3x FLAG peptide (American Peptide, CA) in Low-Salt Buffer. Eluted myosin-15 motor domain was then purified by anion exchange chromatography (5/50 MonoQ GL; Cytiva). After injecting the sample, the column was washed with 10 mM MOPS, 100 mM NaCl, 1 mM EGTA, 0.1 mM PMSF, 1 mM DTT, pH 7.0, then eluted with a linear gradient to 1M NaCl. Fractions eluting at ∼31 mS·cm^−1^ were concentrated (10,000 MWCO) and further purified by size exclusion chromatography (Superdex 200, Cytiva) with isocratic elution in 10 mM MOPS, 100 mM KCl, 0.1 mM EGTA, 1 mm NaN_3_, 0.1 mM PMSF, 1 mM DTT, 1 μg·mL^−1^, pH 7.0. The myosin-15 : ELC : RLC complex (1:1:1) eluted as a single peak, and was concentrated (10,000 MWCO) before determining concentration (A280, ε = 88,020 M^−1^·cm^−1^). Purity was confirmed by SDS-PAGE (4 – 20% TGX, BioRad).

### Actin purification

Chicken skeletal muscle actin was purified as previously described (MacLean-Fletcher and Pollard, 1980) and stored in G-Mg buffer: 2 mM Tris-HCl pH 8.0, 0.5 mM DTT, 0.2 mM MgCl_2_, 0.01% NaN_3_ at 4 °C. F-actin was prepared by mixing 5 μM monomeric actin with KMEI buffer (50 mM KCl, 1 mM MgCl_2_, 1 mM EGTA, 10 mM imidazole pH 7.0) supplemented with 0.01% NP40 substitute (Roche). The mixture was incubated at room temperature for 1 hour to stimulate polymerization, followed by 4 °C overnight.

### Grid preparation

For the F-actin alone sample, 3 μl of F-actin diluted to 0.6 μM in KMEI + NP40 was applied to a glow discharged C-flat 1.2/1.3 holey carbon Au 300 mesh grid (Electron Microscopy Sciences) in a Leica GP plunge freezer operating at 25°C. After 60 s incubation, the grid was blotted from the back with Whatman no. 5 filter paper for 4 s and flash frozen in liquid ethane. To prepare the rigor wild-type and *jordan* mutant myosin-15 bound F-actin samples, 10 U/ml apyrase was added to 6 μM myosin-15 and incubated on ice for 20 minutes to remove nucleotide. 60 s after applying 3 μl of 0.6 μM F-actin to a glow discharged grid, 3 μl of myosin-15 was added and incubated for 60 s. After that 3 μl of the mixture solution was removed and another 3 μl of myosin-15 was applied. After an additional 60 s of incubation, 3 μl of the mixture solution was removed and the grid was blotted for 4 s and flash frozen in liquid ethane. The ADP-Mg state wild-type myosin-15 bound F-actin specimen was prepared identically, except that wild-type myosin-15 was incubated with 5 mM ADP-Mg for 20 minutes prior to grid preparation.

### Cryo-EM data collection

Data were collected on a FEI Titan Krios operating at 300 kV equipped with a Gatan K2-summit detector using super-resolution mode. Exposures were targeted using stage translations with a single exposure per hole using the SerialEM software suite (Mastronarde, 2005), and movies were recorded at a nominal magnification of 29,000 X, corresponding to a calibrated pixel size of 1.03 Å at the specimen level (super-resolution pixel size of 0.515 Å / pixel). Each exposure was fractionated across 40 frames with a total electron dose of 60 e^-^ / Å^2^ (1.5 e^-^ / Å^2^ / frame) and a total exposure time of 10 s, with defocus values ranging from -1.5 to -3.5 μm underfocus.

### Cryo-EM image processing

Image processing was carried out using the Iterative Helical Real Space Refinement (IHRSR) protocol as implemented in the RELION 3.0 pipeline (Egelman, 2007; Zivanov et al., 2018). Unless otherwise noted, all steps were carried out in RELION. Movie stacks were motion corrected, dose weighted and summed with 2 x 2 binning (to a 1.03 Å pixel size) using the MotionCor2 algorithm (Zheng et al., 2017) with 5 x 5 patches as implemented in RELION. Contrast Transfer Function (CTF) estimation was performed with CTFFIND4 (Rohou and Grigorieff, 2015). Filaments were auto-picked and split into overlapping segments with a step size of 81 Å, corresponding to 3 actin protomers, then extracted in a box size of 512 pixels and subjected to two-dimensional (2D) classification. Segments contributing to featureful class-averages were selected and subjected to three-dimensional (3D) classification with 4 classes, initialized with helical parameters of 27 Å rise and -166.7° twist. The cryo-EM maps of F actin alone (EMD-7115) and myosin VI bound F actin (EMD-7116) were low pass filtered to 35 Å to serve as the initial models for F actin alone and myosin-15 decorated F actin datasets, respectively. The best 3D class from each dataset was selected and low-pass filtered to 35 Å to serve as the initial reference for subsequent 3D auto-refinement. Segments contributing to all the 3D classes, which did not substantially vary in features, were included for 3D auto-refinement. After 3D auto-refinement, post-processing was performed for each dataset using a 3D mask with a Z length of 50 % of the box size, resulting in maps with 4.23 Å (actin alone), 4.23 Å (rigor wild-type myosin-15), 6.16 Å (rigor *jordan* myosin-15), and 5.56 Å (ADP wild-type myosin-15) resolution based on the Fourier shell correlation (FSC) 0.143 criterion.

To further improve the resolution, iterative CTF refinement, Bayesian polishing and 3D auto-refinement was carried out by adapting a recently described procedure (Mei et al., 2020). Briefly, CTF refinement was initially performed without beam-tilt estimation, followed by another round of 3D-auto-refinement using the converged map from the last refinement as the initial reference, low-pass filtered to 35 Å. Subsequent post-processing was performed using a 3D mask with a Z length of 30 % of the box size, then a second round of CTF refinement with beam-tilt estimation, Bayesian polishing and 3D auto-refinement using the converged map from the previous refinement low-pass filtered to 35 Å as the initial reference. A final masked refinement was performed for the rigor wild-type reconstruction using the 30 % Z length mask, which modestly improved the resolution: masked refinement did not improve either of the other actomyosin reconstructions. Final post-processing for each dataset was performed with a 30 % Z length mask, leading to final resolution assessments of 2.82 Å (F-actin alone), 2.83 Å (rigor wild-type myosin-15), 3.76 Å (rigor *jordan* mutant myosin-15), and 3.63 Å (ADP wild-type myosin-15) by the FSC 0.143 criterion (Figure S1). Local resolution estimation and filtering was performed using RELION 3.0. Data acquisition and processing details are listed in Table S1.

### Model building and structure refinement

For F-actin alone, a published actin model (PDB: 6BNO) (Gurel et al., 2017) was docked into the EM density by rigid body fitting in UCSF Chimera (Pettersen et al., 2004) and manually adjusted in Coot (Emsley and Cowtan, 2004). In this starting model, the D-loop was modeled in the “Out” conformation. For the actomyosin structures, an initial homology model of myosin-15 was generated with I-TASSER (Zhang, 2008) using the crystal structure of myosin V (PDB:1OE9) (Coureux et al., 2003) as the template. The myosin-15 model and the F-actin alone model were then docked into each cryo-EM density map to generate actomyosin starting models.

Models were then subjected to Rosetta density-guided model rebuilding (DiMaio et al., 2015). For each initial model, 200 models were generated, and the top 10 lowest energy models were manually inspected in Coot. Non-overlapping stretches of amino acids that fit the cryo-EM density best were selected from these individual models then stitched together to build the full model. For F-actin alone and rigor wild-type actomyosin-15, the D-loop of a subset of the 10 lowest-energy models adopted the “In” conformation, which were subsequently used to model this conformer for these states. D-loops for all lowest-energy models in the *jordan* mutant reconstruction and wild-type ADP reconstruction adopted the “In” conformation. The stitched full models were refined using phenix.real_space.refine in the Phenix software package (Afonine et al., 2018), iterated with manual adjustment in Coot. Model validation was conducted with MolProbity (Williams et al., 2018) as implemented in Phenix.

### Molecular graphics and multiple sequence analysis

Figures and movies were generated with PyMol (The PyMOL Molecular Graphics System, Version 2.3.4; Schrödinger, LLC), UCSF Chimera (Pettersen et al., 2004), and ChimeraX (Goddard et al., 2018). Multiple sequence alignment was performed with Clustal Omega (Madeira et al., 2019) and formatted with Jalview (Waterhouse et al., 2009).

### Pyrene actin polymerization

Skeletal rabbit actin was purified and labeled on Cys 374 with N-(1-pyrene)-iodoacetamide (Criddle et al., 1985). A correction factor was applied for determining pyrene actin concentration, A_corr_ = A_290_ –(0.127 * A_344_). Actin polymerization was measured in a cuvette-based fluorometer (PTI Quantamaster 400, HORIBA Scientific) with excitation at 365 nm and emission at 407 nm. Gel filtered G-actin monomers (10 % pyrene labeled) were desalted (PD SpinTrap G-25, Cytiva) into a modified G-buffer without ATP (2 mM Tris-HCl, 0.1 mM CaCl_2_, 1 mM NaN_3_, 1 mM DTT, pH 8.0), stored on ice and used within 3 hours. G-actin was converted to the Mg^2+^ bound form by addition of 50 µM MgCl_2_ and 0.2 mM EGTA for two minutes at room temperature, before starting the polymerization reaction by mixing G-actin (3 x stock) with KMEI buffer (1.5x stock) in a 1:2 ratio, respectively. Wild-type myosin-15 and ADP were included in the 1.5 x KMEI buffer, as needed. Final reaction conditions were 2 µM G-actin, 1 µM myosin-15, 50 mM KCl, 1 mM MgCl_2_, 1 mM EGTA, 10 mM imidazole, pH 7.0 at 25 ± 0.1 °C. Data were corrected for the reaction dead-time.

### Single-filament actin polymerization

Skeletal rabbit actin was purified, labeled on Cys 374 using tetramethylrhodamine-5-maleimide (Adipogen Life Sciences), and its concentration measured using A_corr_ = A_290_ –(0.208 * A_550_) (Fujiwara et al., 2002). Biotinylated skeletal muscle actin (#8109-01, HyperMol, Germany) was dialyzed against G-buffer and cleared by ultracentrifugation at 100,000 x *g* for 1 hour. G-actin stocks were prepared with TMR-actin (20 %) and biotin-actin (10 %) doping, and desalted into a modified ATP-free G-buffer as described above. Cover glass (24 × 50 mm, #1.5, Marienfeld Superior, Germany) were sequentially sonicated in 2% Hellmanex III (Hellma, Germany), 1 M KOH, 100 % ethanol, and finally Milli-Q water, prior to drying under a nitrogen stream and processing under argon plasma (ZEPTO, Diener Electronic, Germany). Cleaned cover glass were coated with mPEG-silane (2 mg / mL, Laysan Bio, AL) and biotin–PEG–silane (2 μg·mL^-1^, Laysan Bio) diluted in 96 % ethanol, 0.1 % (v/v) HCl, and baked at 70 °C for 1 hour. Cover slips were rinsed in 96 % ethanol, sonicated, followed by rinsing and sonication in Milli-Q water, and finally dried under a filtered nitrogen stream. Flow chambers were assembled using double-sided sticky tape to create a channel on a glass slide. Flow cells were washed with T50 buffer (10 mM Tris·HCl, 50 mM KCl, 1 mM DTT, pH 8.0), and incubated with 0.1 mg / mL neutravidin (Thermo Scientific) in T50 for 1 minute, then washed in 1 mg / mL bovine serum albumin (A0281, Sigma Aldrich) in T50 for 30 s, followed by a final wash of T50.

Experiments were performed in the following buffer: 50 mM KCl, 1 mM MgCl_2_, 1 mM EGTA, 10 mM imidazole, 10 mM DTT, 15 mM glucose, 0.5 % methylcellulose, 20 μg·mL^-1^ catalase, 100 μg·mL^- 1^ glucose oxidase, pH 7.0 at 21 ± 1 °C. Purified myosin-15 motor domain (1 µM) and ADP were included as needed. The reaction flow cell was imaged using an inverted microscope (Nikon Ti-E2) equipped with an oil immersion objective (CFI Apochromat TIRF 100x, 1.49 N.A., Nikon) for objective-style total internal reflection fluorescence (TIRF) microscopy (H-TIRF, Nikon). TMR fluorescence was excited using 561 nm, and emission filtered (ET630/75m, Chroma). Images were captured on an EM-CCD camera (iXon Ultra 888, Andor) controlled by NIS-Elements (AR version 5.2, Nikon). Data were corrected for the assay dead-time. Images were background subtracted and registered (descriptor-based series registration, 2d/3d +t) in FIJI (https://fiji.sc). Actin filament densities were quantified using the Analyze Particle command (size > 3 pixel^2^, circularity: 0.0-1.0) to count particles within a 50 x 50 μm region of interest (ROI) that was randomly selected from the image. A minimum of 3 experiments, from two independent protein preparations, were analyzed for each condition. Filament elongation rates were calculated using kymographs generated in Elements Software (Nikon).

## Notes

### Competing Interest Statement

The authors have declared no competing interest.

### Summary of Updates

We have updated the citation to our companion paper (Moreland et al. 2021) which has now also been posted as a bioRxiv pre-print: doi: 2021.07.09.451618 This version is otherwise identical.

